# m^6^A regulates the stability of cellular transcripts required for efficient KSHV lytic replication

**DOI:** 10.1101/2023.02.09.527825

**Authors:** Oliver Manners, Belinda Baquero-Perez, Timothy J. Mottram, Ivaylo D. Yonchev, Christopher J. Trevelyan, Molly R. Patterson, Andrew Macdonald, Stuart A. Wilson, Julie L. Aspden, Adrian Whitehouse

## Abstract

The epitranscriptomic modification *N*^6^-methyladenosine (m^6^A) is a ubiquitous feature of the mammalian transcriptome. It modulates mRNA fate and dynamics to exert regulatory control over numerous cellular processes and disease pathways, including viral infection. Kaposi’s sarcoma-associated herpesvirus (KSHV) reactivation from the latent phase leads to redistribution of m^6^A topology upon both viral and cellular mRNAs within infected cells. Here we investigate the role of m^6^A in cellular transcripts upregulated during KSHV lytic replication. Results show that m^6^A is crucial for the stability of the *GPRC5A* mRNA, whose expression is induced by the KSHV latent-lytic switch master regulator, the replication and transcription activator (RTA) protein. Moreover, we demonstrate that GPRC5A is essential for efficient KSHV lytic replication by directly regulating NFκB signalling. Overall, this work highlights the central importance of m^6^A in modulating cellular gene expression to influence viral infection.

**Author Summary:** Chemical modifications on mRNA, such as m^6^A, are functionally linked to all stages of mRNA metabolism and regulate a variety of biological processes. As such, m^6^A modification offers unique possibilities for viruses to modulate both viral and host gene expression. m^6^A has been identified on transcripts encoded by a wide range of viruses and studies to investigate m^6^A function have highlighted distinct roles in virus life cycles. In addition, cellular transcripts undergoing differential m^6^A status during infection may also be important for virus replication. In this study we investigate the impact of differential m^6^A modification in host transcripts during KSHV lytic replication, by identifying transcripts with altered methylation profiles between latent and lytic replication programmes. We show that increased m^6^A content in one of these cellular mRNAs, *GPRC5A*, enhances its stability and correlates with increased abundance during KSHV lytic replication. Moreover, the importance of GPRC5A is demonstrated by depletion studies, showing that GPRC5A enhances KSHV lytic replication by inhibiting cell signalling pathways.

## Introduction

Cellular infection by viruses, including those in the *Herpesviridae*, is greatly influenced by post-transcriptional regulatory mechanisms which modulate gene expression. Kaposi’s sarcoma-associated herpesvirus (KSHV) is the aetiological agent of Kaposi’s sarcoma and the B cell lymphoproliferative disorders, primary effusion lymphoma and multicentric Castleman’s disease [1–4]. KSHV undergoes a complex biphasic life cycle encompassing latent persistence and lytic replication phases. During latency, the virus enters a state of transcriptional dormancy where expression is limited to a few latent genes [5–7]. Certain factors, including stress or immunosuppression, trigger the reactivation of the virus into the lytic phase where over 80 viral genes are expressed leading to the production of infectious virions [8, 9]. The replication and transcription activator (RTA) protein, expressed from the open reading frame ORF50, is essential and sufficient for the process of reactivation and serves as a molecular switch from latent to lytic replication phases [10–12]. Notably, both the latent and lytic replication phases are essential for KSHV-mediated tumorigenicity [13–15].

Amassing evidence suggests that viral life cycles are greatly influenced by a group of chemical modifications of RNA, collectively known as the epitranscriptome [16]. The most common epitranscriptomic modification of mRNA is N^6^-methyladenosine (m^6^A), whose dynamics are controlled by specific cellular machinery which install, remove and decode the modification [17–20]. The multi-component m^6^A writer complex comprises the catalytically active subunit, methyltransferase-like 3 (METTL3) and methyltransferase-like 4 (METTL14) which methylates the central adenosine residue at the consensus DRACH sequence (D=A/G/U, R=A/G and H=A/C/U) [12]. Additional components, such as WTAP, RBM15, VIRMA and ZC3H13 are responsible for selectivity, localisation and structural integrity [21–23]. Conversely, two RNA demethylases, known as m^6^A erasers, α-ketoglutarate-dependent dioxygenase alkB homolog 5 (ALKBH5) [24] and fat mass obesity protein (FTO) [25] revert m^6^A back to adenosine residues. Although the impact and extent of demethylation carried out by the m^6^A ‘erasers’ ALKBH5 and FTO is debated, it is widely accepted that at least some m^6^A residues can be reversed back to adenosine [26]. Importantly, the dynamic reversible addition and removal of m^6^A allows rapid adjustment of mRNA fate and thus regulatory control. Finally, m^6^A exerts its influence over mRNAs *in cis* by recruiting RNA binding proteins, known as m^6^A readers, which in turn modulate the structure, splicing, nuclear export, stability and translation of the transcript [27]. The most widely characterised group of m^6^A readers, the YT521-B homology (YTH) domain containing proteins, directly bind m^6^A in target RNAs through an aromatic cage [28]. In contrast, a second group of m^6^A readers, including several hnRNPs, preferentially bind m^6^A-modified RNAs through an m^6^A switch mechanism, an event in which m^6^A modification remodels local RNA structure [29]. A myriad of m^6^A readers may therefore exist which enable widespread regulatory control over gene expression [19], affecting many biological pathways. Importantly, dysregulation of these pathways is now implicated in a wide variety of disease states, including viral infection.

The ability of m^6^A to dynamically regulate gene expression offers unique possibilities for viruses to modulate viral and host gene expression, but also for the host to regulate a response to infection [30]. Recent studies demonstrate that m^6^A is deposited upon the RNAs transcribed by a diverse range of both RNA and DNA viruses [31–34], and highlighted distinct pro and antiviral roles indicating widespread regulatory control over viral life cycles [16, 34]. However, whilst the modification of viral RNAs has been investigated in multiple studies, the impact of m^6^A sites upon cellular transcripts during viral infection largely remains unknown [31, 32, 35]. Importantly, the redistribution of m^6^A topology in response to viral infection has been described, but very few studies have reported on the impact of cellular m^6^A on a site-specific basis. Therefore, determining the impact of individual m^6^A residues in cellular mRNAs during viral infection is a key step in the advancement of viral epitranscriptomics.

In this study we investigate the impact of m^6^A in host transcripts upon KSHV reactivation, by identifying transcripts with altered methylation profiles between latent and lytic replication programmes. We identify a subset of cellular mRNAs which heavily increase in both m^6^A content and abundance during KSHV lytic replication. Functional interrogation suggests these genes encode key cellular factors involved in enhancing lytic replication and whose expression can be reduced through the disruption of m^6^A sites in these transcripts, indicating these m^6^A sites have potential for therapeutic targeting.

## Results

### Differential m^6^A status of host cell transcripts corresponds to expression levels during KSHV lytic replication

m^6^A-seq analysis obtained from KSHV-infected TREx BCBL1-Rta cells, a latently infected KSHV B-lymphocyte cell line that expresses a Myc-tagged viral RTA under the control of a doxycycline-inducible promoter, identified large scale remodelling of the host cell m^6^A epitranscriptome during reactivation of the KSHV lytic replication cycle [32]. However, given that many of these changes in m^6^A peaks were concordant with changes in mRNA expression, we specifically searched for mRNAs with differentially modified m^6^A sites relative to an m^6^A peak in the same transcript whose methylation status remained stable. Notably, we identified 58 transcripts which fulfilled such criteria (**S1 Table**) and examples of differential m^6^A modification within specific transcripts during latent and lytic replication phases are demonstrated in **S1 Fig**. Ingenuity pathway analysis (IPA) was then performed to assess whether these differentially m^6^A modified mRNAs were associated with distinct functional networks and cellular activities (**S2 Fig**.). The strongest predicted canonical pathways involved RNA processing and cell signalling. Similarly, predicted molecular and cellular functions and diseases identified cell-to-cell signalling, gene expression and cancer-related pathways.

We then prioritised cellular mRNAs which increased in their expression during lytic replication, as these contrast with the majority of cellular transcripts which are degraded by the viral SOX endonuclease during KSHV-mediated host cell shutoff [36, 37]. To investigate changes in host mRNAs during lytic replication, we selected a number of transcripts with the greatest changes in m^6^A content and compared their expression levels during latency and 24 hours post-induction of lytic replication by RT-qPCR (**Fig. 1A**). The selected mRNAs were significantly upregulated in the lytic phase compared with latency. Most notably, *GPRC5A* and *FOSB* transcripts were 19-fold more abundant in the lytic phase. Immunoblotting of cell lysates from latent and lytic TREx BCBL1-Rta cells using a GPRC5A-specific antibody confirmed the changes in mRNA expression led to a similar increase in protein levels (**Fig. 1B**).

**Figure 1.**
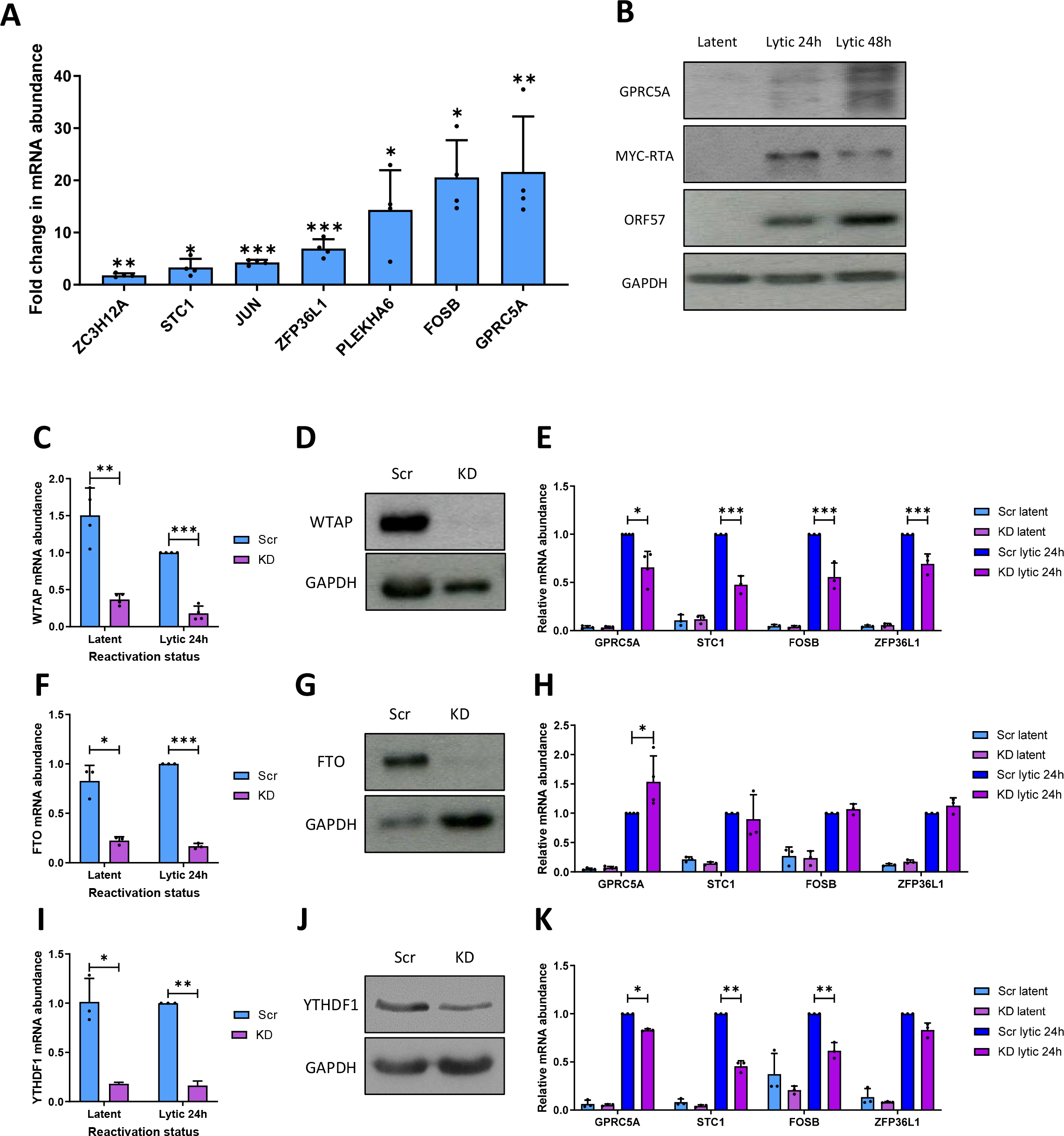
m^6^A affects the abundance of host transcripts upregulated during KSHV reactivation. (A) RT-qPCR analysis of cellular m^6^A-modified transcripts in TREx BCBL1-Rta cells induced for 24 hours compared to latent levels (mean ±SD, n=4). (B) Representative western blot of STC1 and GPRC5A protein levels in latent and reactivated TREx BCBL1-Rta cells (blots are representative of 3 biological replicates). (C) RT-qPCR analysis of *WTAP* RNA levels in latent and lytic WTAP shRNA-treated TREx BCBL1-Rta cells (mean ±SD, n=3). (D) Representative western blot of WTAP protein levels in Scr and WTAP-shRNA treated latent TREx BCBL1-Rta cells. (E) RNA levels of *GPRC5A, STC1, FOSB* and *ZFP36L1* in Scr- or WTAP shRNA-treated latent and induced TREx BCBL1-Rta cells (mean ± SD, n=3). (F) RT-qPCR analysis of *FTO* RNA levels in latent and lytic FTO shRNA-treated TREx BCBL1-Rta cells (mean ±SD, n=3). (G) Representative western blot of FTO protein levels in Scr and FTO-shRNA treated latent TREx BCBL1-Rta cells. (H) RNA levels of *GPRC5A, STC1, FOSB* and *ZFP36L1* in Scr- or FTO shRNA-treated latent and induced TREx BCBL1-Rta cells (mean ± SD, n=3). (I) RT-qPCR analysis of *YTHDF1* RNA levels in latent and lytic YTHDF1 shRNA-treated TREx BCBL1-Rta cells (mean ±SD, n=3). (J) Representative western blot of YTHDF1 protein levels in Scr and YTHDF1-shRNA treated latent TREx BCBL1-Rta cells. (K) RNA levels of *GPRC5A, STC1, FOSB* and *ZFP36L1* in Scr- or YTHDF1 shRNA-treated latent and induced TREx BCBL1-Rta cells (mean ± SD, n=3).

Given a potential link between the increase in cellular transcript abundance and increased m^6^A content during KSHV lytic reactivation (**S1 Table**), we speculated that disruption of m^6^A dynamics, through depletion of the host cell m^6^A machinery components, may alter mRNA levels. To examine this hypothesis, TREx BCBL1-Rta cells were stably transduced with lentivirus-based shRNAs targeting the m^6^A writer (WTAP), the m^6^A eraser (FTO) or the m^6^A reader (YTHDF1). RT-qPCR and immunoblotting confirmed the reduction of these proteins by 75% (WTAP), 73% (FTO) and 82% (YTHDF1) (**Fig. 1C, D, F, G, I and J**). Host cell mRNA levels were then examined following disruption of the m^6^A machinery during latency or lytic replication (**Fig. 1E, H and K**). Notably, WTAP depletion significantly reduced the abundance of the cellular m^6^A modified mRNAs: *GPRC5A, STC1, FOSB* and *ZFP36L1*, suggesting that methylation is important for their abundance during KSHV lytic replication. Conversely, the depletion of the m^6^A eraser FTO enhanced the upregulation of *GPRC5A* by ~50% (**Fig. 1K**). However, we did not detect a significant increase in the levels of *STC1, FOSB* or *ZFP36L1* suggesting that FTO may not target these transcripts for demethylation. Finally, knockdown of the m^6^A reader YTHDF1 recapitulated the effect of WTAP depletion, with diminished levels of *GPRC5A, STC1, FOSB* and *ZFP36L1* observed. Taken together, these data suggest that m^6^A is linked to the abundance and upregulation of four m^6^A-modified cellular mRNAs; *GPRC5A, STC1, FOSB* and *ZFP36L1*.

### m^6^A sites within *GPRC5A* mRNA regulates its stability

Although depletion of WTAP, FTO and YTHDF1 and the consequent disruption of m^6^A dynamics modulated *GPRC5A, STC1, FOSB* and *ZFP36L1* transcript levels, it remained unclear whether these changes are direct *cis* acting effects on these mRNAs resulting from perturbation of the m^6^A machinery. Therefore we used m^6^A immunoprecipitation coupled with RT-qPCR (m^6^A IP-qPCR) to validate the presence of m^6^A methylation on these cellular transcripts (**Fig. 2A**). Transcripts *SLC39A14* and *GAPDH* were used as positive and negative controls, as the former contains a prominent and well described m^6^A site, while the latter is known to be an unmodified transcript [18]. For each gene, two primer sets were designed with one spanning the putative m^6^A peak and the other spanning a region lacking m^6^A according to the m^6^A-seq data. m^6^A peaks were confirmed by enrichment of the m^6^A spanning over the null primer sets by m^6^A IP-qPCR using RNA fragmented to ~200nt to ensure good spatial resolution. Prominent m^6^A modification was detected in *SLC39A14, GPRC5A, FOSB* and *ZFP36L1* mRNAs, while no enrichment was identified for *GAPDH*, validating the m^6^A modification of *GPRC5A, FOSB* and *ZFP36L1* at the expected locations. Unfortunately, this method could not be used for *STC1* as no suitable primer sets could be designed which were able to amplify the proposed methylated region efficiently.

**Figure 2.**
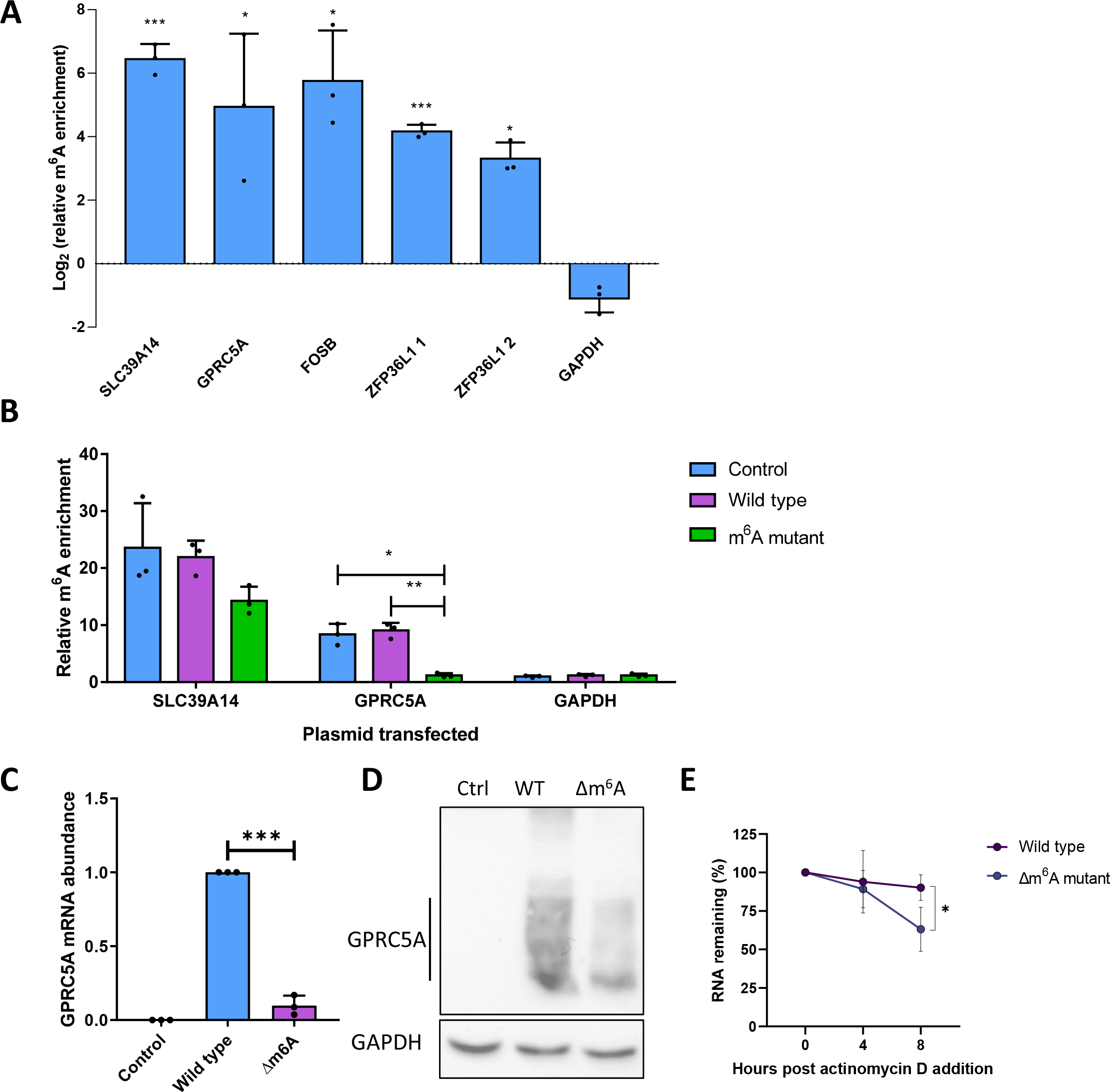
m^6^A sites within the cellular *GPRC5A* transcript are essential for its stability. (A) m^6^A IP-qPCR analysis of cellular transcripts or control RNAs in latent TREx BCBL1-Rta cells showing log2 fold enrichment of m^6^A-modifed region relative to a non-methylated region in cis. 2 regions were analysed within ZFP36L1 as this mRNA contains a wide flat m^6^A peak spanning several hundred bp (mean ± SD, n=3). (B) m^6^A IP-qPCR analysis of *GPRC5A* or control RNAs in GPRC5A wild type or Δm^6^A mutant-transfected HEK 293T cells showing fold enrichment of m^6^A-modifed region relative to a non-methylated region in cis (mean ± SD, n=3). (C) RT-qPCR analysis of *GPRC5A* RNA in cells transfected with control plasmid, wild type of GPRC5A-expressing constructs (mean ± SD, n=3). (D) Representative western blot of GPRC5A protein levels in cells transfected with no plasmid, wild type or Δm^6^A GPRC5A-expressing constructs. (E) RT-qPCR analysis of wild type or Δm^6^A *GPRC5A* RNA in cells treated with actinomycin D for 0, 4 or 8 hours (mean ± SD, n=3).

We prioritised GPRC5A for further investigation as it had a single narrow differentially modified m^6^A peak and its abundance was altered during lytic replication of TREX-BCBL1-RTA cells depleted for WTAP, FTO or YTHDF1. The 4 potential DRACH sites within the *GPRC5A* m^6^A peak were altered by site-directed mutagenesis, termed Δm^6^A, with no alteration in amino acid sequence and minimal changes to the DNA sequence (**S3 Fig**.). m^6^A IP-qPCR confirmed that the Δm^6^A mRNA was devoid of the modification, unlike both endogenous and wild type transfected mRNAs (**Fig. 2B**). Furthermore, deletion of m^6^A within GPRC5A recapitulated the effects of WTAP or YTHDF1 depletion, with a reduction of *GPRC5A* mRNA and protein levels (**Fig. 2C and D**). To assess whether the reduced mRNA levels were due to decreased stability, cells were transfected with wild type or Δm^6^A GPRC5A constructs and treated with the transcriptional inhibitor actinomycin D for 0, 4 or 8 hours. RT-qPCR analysis showed a significant reduction in stability in the Δm^6^A mutant, resulting in only 63% of Δm^6^A *GPRC5A* after 8 hours of actinomycin D treatment compared with 90% for the wild type construct (**Fig. 2E**). Taken together, these results suggest a differentially modified m^6^A peak, containing 4 potential DRACH sites, within the *GPRC5A* mRNA is important for its abundance by enhancing its stability.

### RTA transactivates the GPRC5A promoter

Cellular transcripts which increase in abundance during KSHV lytic replication contrast with the majority of host mRNAs which are degraded by the KSHV-encoded endonuclease, SOX [36, 37]. As a result, upregulated transcripts are more likely to be functionally relevant in the KSHV lytic replication cycle. Two KSHV lytic proteins with the ability to increase the abundance of host transcripts during the early stages of lytic replication, are the lytic master regulator RTA, encoded by ORF50, and the mRNA processing factor ORF57. RTA is a transcription factor able to transactivate both viral and cellular promoters [38], whereas ORF57 can bind and stabilise cellular mRNAs, enhancing their RNA processing and translation [39]. To determine if the upregulation of the m^6^A-modified *GPRC5A* transcript is dependent on these viral factors, we transfected constructs expressing GFP-tagged-RTA [40] or GFP-tagged-ORF57 [41] into HEK 293T cells and confirmed their ectopic expression by western blotting (**Fig. 3A**). Results showed a 3-fold increase in *GPRC5A* mRNA, but only in the presence of GFP-tagged-RTA expression (**Fig. 3B**). Furthermore, by transfecting increasing amounts of GFP-tagged-RTA into HEK 293T cells, we identified a dose-dependent increase in *GPRC5A* mRNA levels up to 4-fold (**Fig. 3C**). Additionally, the increase in *GPRC5A* mRNA was accompanied by a 6-fold increase in GPRC5A protein production (**Fig. 3D and E**). This is supported by previous genome-wide RTA ChIP-seq studies which identified RTA binding sites within the GPRC5A promoter (**S4 Fig**.) [42]. Together, these results suggest that the KSHV protein RTA induces the transcription of *GPRC5A* mRNA, whose posttranscriptional stability is regulated by m^6^A modification.

**Figure 3.**
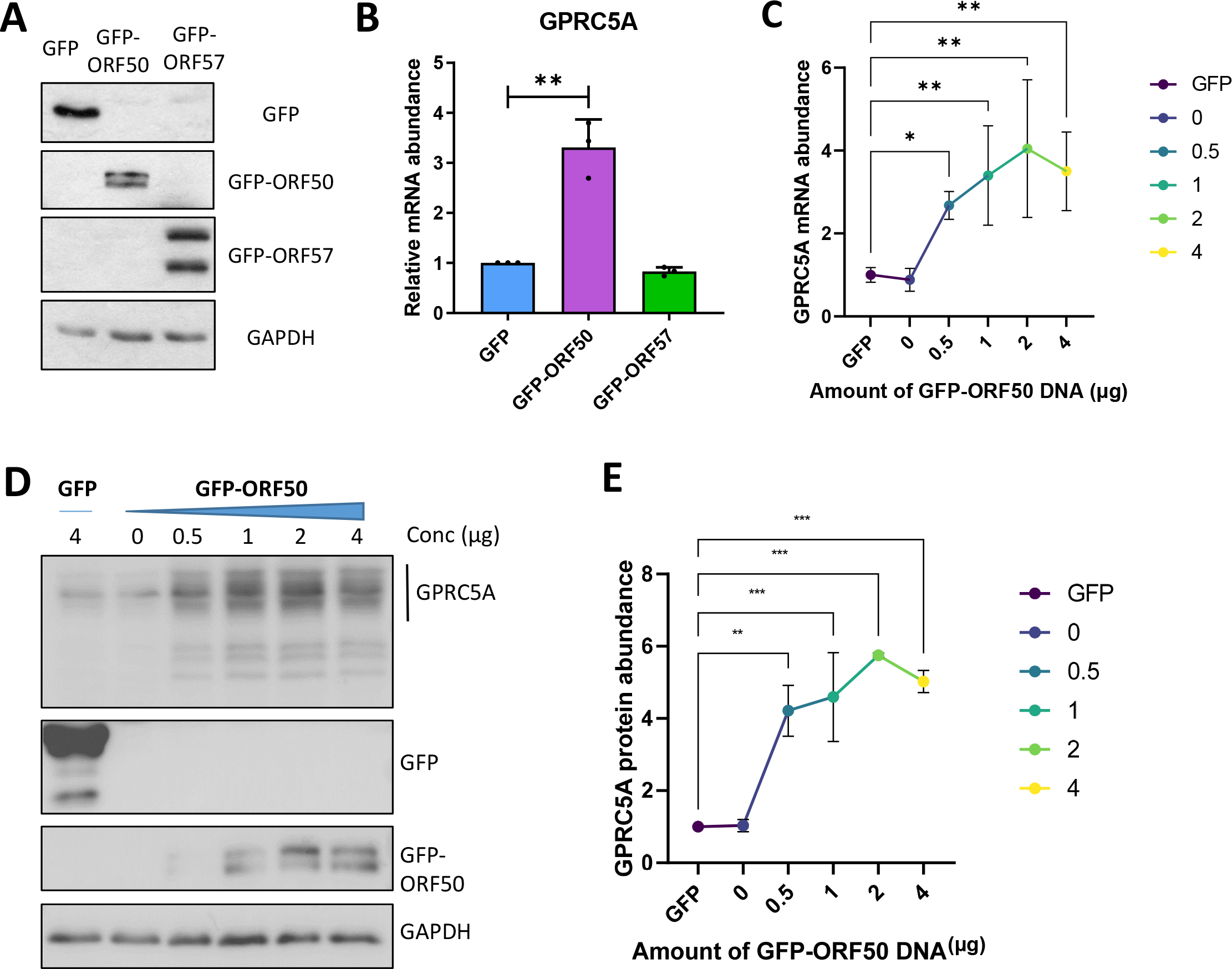
GPRC5A is induced by the KSHV lytic transactivator, RTA. (A) Representative western blot showing expression of transfected GFP-tagged viral proteins ORF50 and ORF57 in naive uninfected 293T cells. (B) RT-qPCR analysis of *GPRC5A* RNA levels in viral protein transfected 293T cells (mean ± SD, n=3). (C) RT-qPCR analysis of *GPRC5A* RNA levels in cells transfected with increasing concentrations of GFP-ORF50 DNA (mean ± SD, n=3). (D) Representative western blot of GPRC5A protein levels in cells transfected with a range of GFP-ORF50 DNA concentrations. (E) Densitometry quantification of immunoblots was performed using the Image J software and is shown as a percentage relative to the loading control, GAPDH (mean ±SD, n=3).

### GPRC5A is required for efficient KSHV lytic reactivation

Given that expression of GPRC5A is strongly induced by RTA during KSHV lytic reactivation and its transcript is additionally stabilised by m^6^A modification, we hypothesised that GPRCA5 may be important for KSHV lytic replication. To investigate this hypothesis, GPRC5A was depleted from TREx BCBL1-Rta cells by lentiviral transduction expressing two alternative GPRC5A-targeting shRNAs and knockdown was confirmed by RT-qPCR (**Fig. 4A**). During latency, a 40% and 60% decrease in *GPRC5A* mRNA levels were achieved relative to cells treated with a scrambled control shRNA. Moreover, both shRNAs inhibited the upregulation of *GPRC5A* mRNA during reactivation, resulting in a more significant reduction of 80% during lytic replication. Upon reactivation of control and GPRC5A knockdown TREx BCBL1-Rta cells, a slight but significant 18% reduction in the levels of the early lytic *ORF57* mRNA with a greater 29-43% decrease in the late lytic *ORF47* transcript was observed, suggesting cumulative inhibition of the KSHV lytic cycle (**Fig. 4B**). Similarly, reduced ORF57 (early lytic) and ORF65 (late lytic) protein expression was identified in GPRC5A-depleted cells (**Fig. 4C and D**). Furthermore, reinfection assays were performed using supernatant harvested from scrambled and GPRC5A-depleted TREx BCBL1-Rta cells to reinfect naïve HEK 293T cells (**Fig. 4E**). Measuring *ORF57* mRNA levels in these reinfected HEK 293T cells indicated that knockdown of GPRC5A lead to a significant 38-56% decline in the production of infectious virus particles relative to cells treated with a scrambled shRNA. Together, these experiments suggest that GPRC5A is upregulated during KSHV reactivation to enhance lytic replication and efficient virion production.

**Figure 4.**
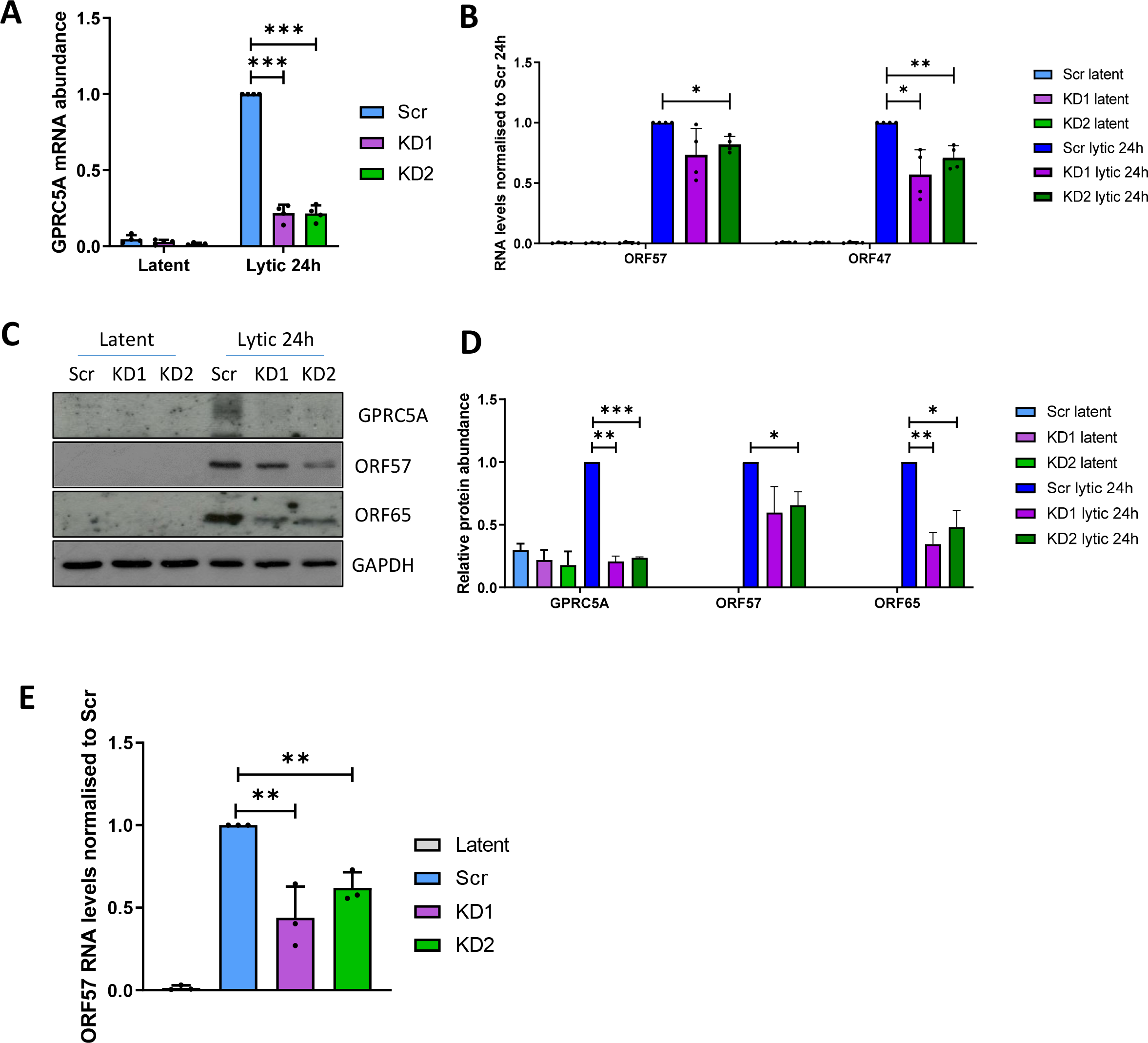
GPRC5A depletion reduces KSHV lytic replication. (A) RT-qPCR analysis of *GPRC5A* RNA levels in latent and lytic GPRC5A shRNA-treated TREx BCBL1-Rta cells (mean ±SD, n=3). (B) RT-qPCR analysis of viral *ORF57* and *ORF47* RNAs in latent and lytic GPRC5A-depleted cells (mean ±SD, n=3). (C) Representative western blot of viral proteins ORF57 and ORF65 in latent and lytic GPRC5A-depleted cells. (D) Densitometry quantification of immunoblots was performed using the Image J software and is shown as a percentage relative to the loading control, GAPDH (mean ±SD, n=3). (E) RT-qPCR of *ORF57* RNA levels in 293T cells treated with virus-containing supernatant harvested from scrambled or GPRC5A shRNA-treated TREx BCBL1-Rta cells (mean ±SD, n=3).

### GPRC5A inhibits NFκB signalling to support KSHV lytic replication

Multiple studies have reported a role for membrane bound G-protein coupled receptors (GPCRs) at the top of signalling cascades, although their contribution to viral replication has not been thoroughly explored [43]. Interestingly, KSHV encodes a constitutively active GPCR (vGPCR) during lytic replication, which participates in oncogenic signalling through the ERK1/MAPK pathway to induce angiogenesis. To investigate why KSHV also upregulated host cell GPRC5A during lytic replication we mapped its viral and cellular cofactors. To identify such interactors, TREx BCBL1-Rta cells were transduced with a constitutive expressing GPRC5A-FLAG expressing construct and global quantitative proteomic analysis was performed on FLAG immunoprecipitates from latent and reactivated TREx BCBL1-Rta cell lysates (**Fig. 5A**). We identified 16 proteins with greater than 1.5-fold interaction with GPRC5A during lytic replication (**S2 Table**). Among these proteins, both members of the flotillin family, FLOT1 and FLOT2, were present at 4-fold and 27-fold higher levels in the lytic phase, respectively (**Fig. 5B**). This result was confirmed by coimmunoprecipitation of FLOT1 in GPRC5A-GFP immunoprecipitates from latent and lytic TREx BCBL1-Rta cells, where increased FLOT1 protein was detected in association with GPRC5A during KSHV lytic replication (**Fig. 5C**). Furthermore, immunofluorescence studies revealed that FLOT1 and GPRC5A-GFP were both concentrated at the cell periphery in membrane-associated microdomains reminiscent of lipid rafts in both replication states; with enhanced co-localisation observed during lytic replication (**Fig. 5D**). As GPCRs are commonly associated with cellular signalling pathways we hypothesised that improved organisation of GPRC5A into these membrane-associated microdomains during lytic replication may allow for increased signal transduction through this protein [44].

**Figure 5.**
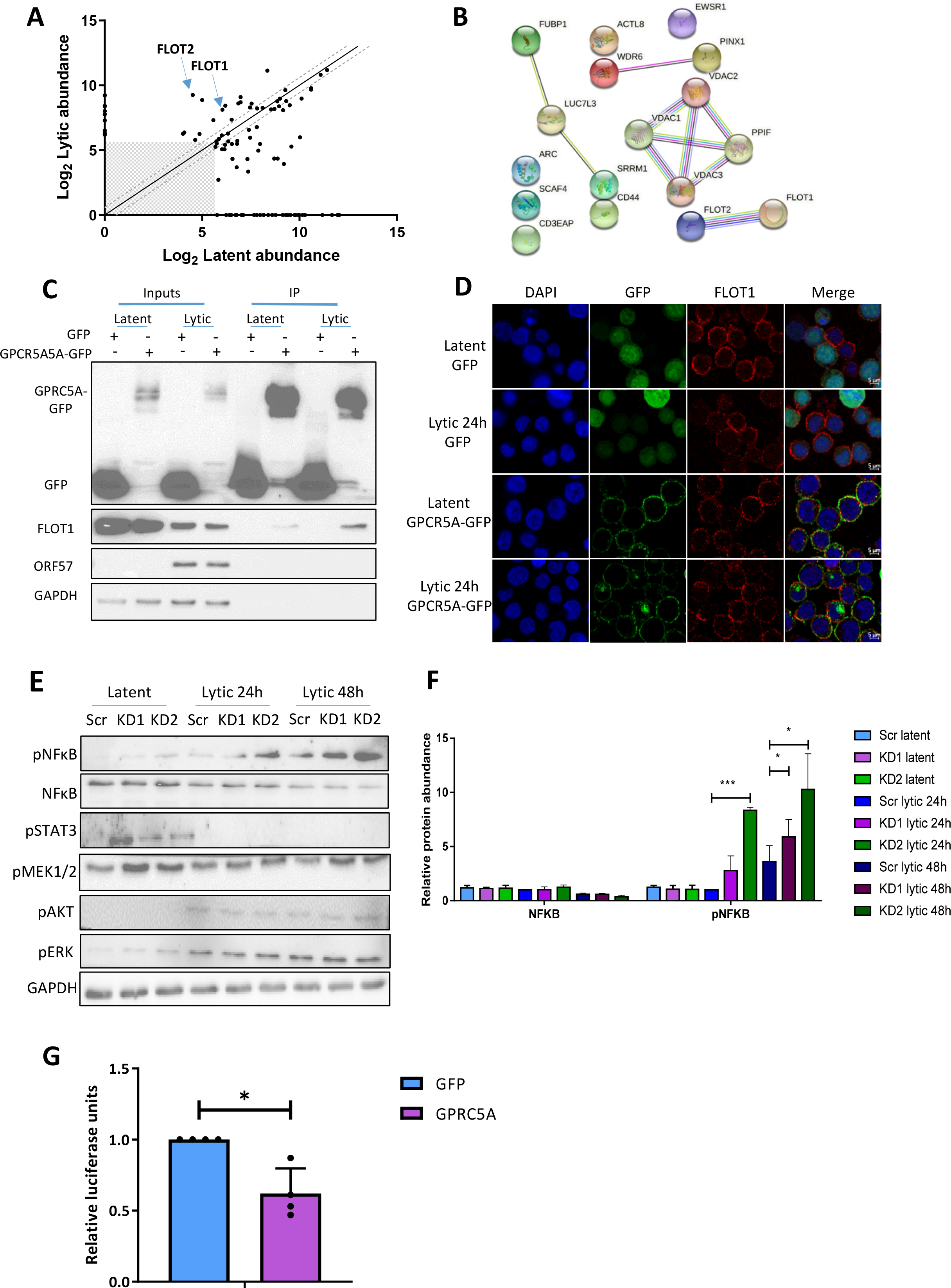
GPRC5A interacts with members of the Flotillin family and affects NFκB signalling. (A) Proteins with >1.5 enrichment in FLAG-immunoprecipitates from GPRC5A-FLAG-overexpressing TREx BCBL1-Rta cells compared with parental TREx BCBL1-Rta cells after TMT-mass spec analysis. (Grey box: data points with less than 50 abundance in both latent and lytic TREx BCBL1-Rta cells were excluded). (B) STRING interaction map showing proteins with >1.4 fold interaction with GPRC5A in lytic replication compared with latency in TREx BCBL1-Rta cells after TMT-mass spec analysis. (C) Immunoprecipitation of GFP or GPRC5A-GFP from transduced latent and lytic TREx BCBL1-Rta cells showing increased co-immunoprecipitation of FLOT1 in lytic replication. (D) Immunofluorescence analysis of FLOT1 localisation in latent and lytic TREx BCBL1-Rta cells transduced with a GFP control (upper two) or GFP-tagged GPRC5A (lower two) expressing plasmid. (E) Representative western blots of NFκB, pNFκB, pSTAT, pMEK1/2, pAKT, pERK and GAPDH protein expression in latent and lytic GPRC5A-depleted cells. (F) Densitometry quantification of immunoblots was performed using the Image J software and is shown as a percentage relative to the loading control, GAPDH (mean±SD, n=3). (G) Luciferase reporter assay from HEK-293Ts co-transfected with GFP or GPRC5A-GFP alongside various described signalling reporters with transcription factor binding sites attached to a luciferase reporter plasmid. Data presented are relative to an internal firefly control (mean ±SD, n=3).

Given the widespread role of GPCRs in cellular signalling pathways, many of which are of central importance within KSHV lytic replication, we sought to identify whether upregulation of GPRC5A affects downstream signalling cascades which may impact KSHV lytic replication. Through western blot analysis of key signalling proteins comparing scrambled control and GPRC5A-depleted TREx BCBL1-Rta cells undergoing lytic replication, we observed little to no change in pERK, pAKT, pMEK1/2 or pSTAT3 levels. However, results identified a significant increase in levels of p-NFĸB S536 (p65) in cells depleted for GPRC5A compared to those treated with a scrambled control, indicating an increase in NFĸB signalling (**Fig. 5E and F**). To support this observation and determine whether GPRC5A directly regulates NFκB signalling, we transfected HEK 293T cells with GPRC5A-GFP or GFP control expression constructs alongside an NFκB-responsive luciferase reporter plasmid (**Fig. 5G**). Analysis showed luciferase activity was significantly attenuated in HEK 293T cells transfected with GPRC5A-GFP compared to those transfected with control GFP, suggesting inhibition of NFĸB activity. These results are consistent with previous observations showing that NFĸB is reduced during KSHV lytic replication [45], and suggest that GPRC5A contributes to this downregulation.

## Discussion

Although m^6^A is known to play a crucial regulatory role in the life cycle of numerous viruses, most studies to date have investigated the contribution of the modification to viral transcripts, rather than cellular gene expression. Here, we show that a number of host cell transcripts with differential m^6^A modification during the latent and lytic replication phases are highly expressed during lytic replication. In addition, focussing on one of these differentially m^6^A modified transcripts, we show that GPRC5A enhances KSHV lytic replication by acting as a novel inhibitor of NFκB signalling (**Fig. 6**).

**Figure 6.**
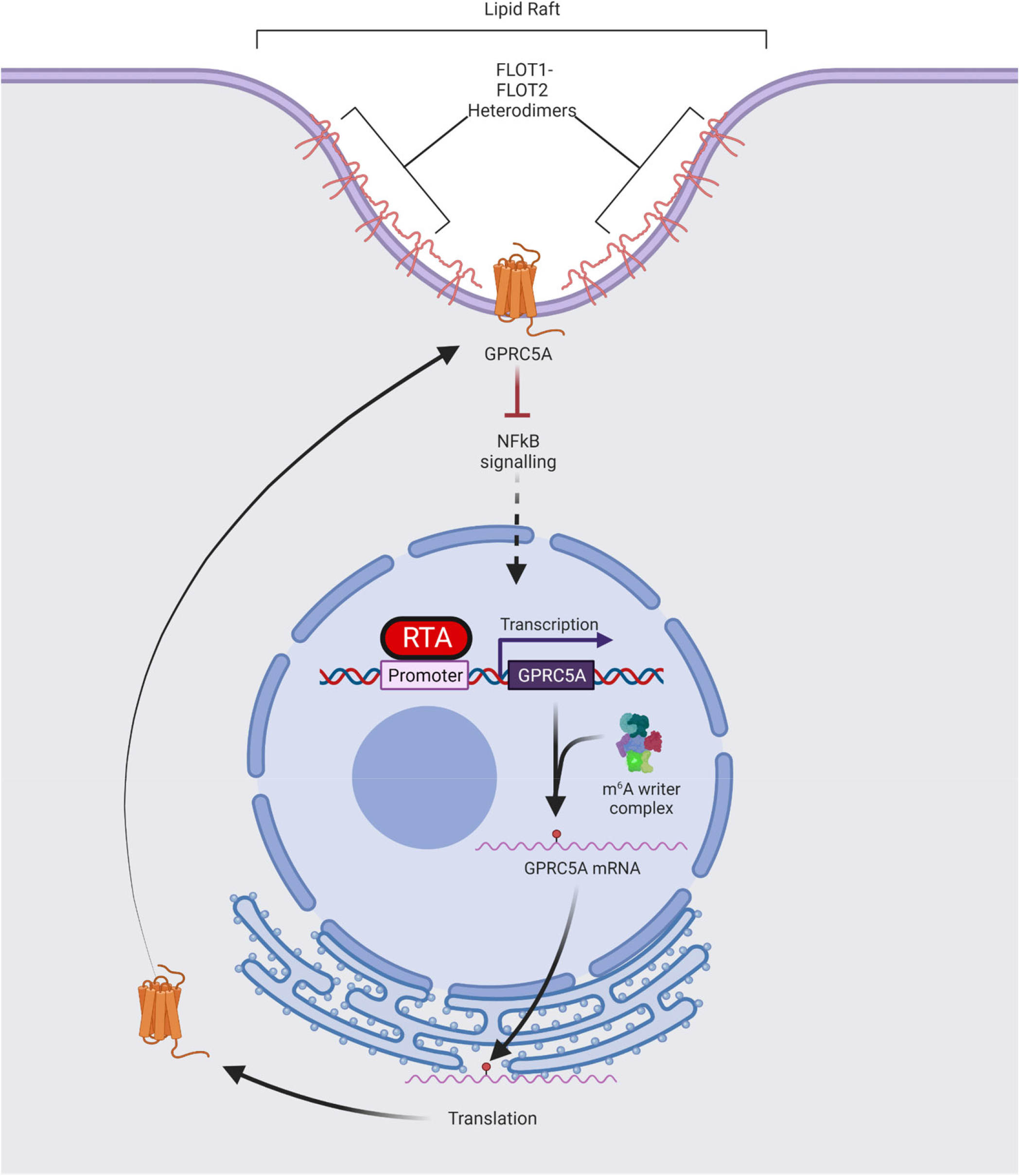
Schematic showing potential regulation of GPRC5A gene expression during KSHV lytic replication. At the onset of KSHV lytic replication, the viral lytic master regulator RTA transactivates the GPRC5A promoter increasing transcription of *GPRC5A* transcripts which then becomes m^6^A-modified by the m^6^A methyltransferase complex. Methylation of *GPRC5A* mRNA increases its stability *in cis* allowing translation of a larger pool of GPRC5A transcripts and therefore enhanced protein expression. Once translated, GPRC5A is transported to the cell membrane where it is organised into lipid rafts composed of members of the flotillin family. As a membrane-bound GPCR, GPRC5A inhibits NFκB signalling to enhance KSHV lytic replication.

Previous research has shown that KSHV lytic replication induces vast changes in m^6^A distribution with important consequences for both host and viral gene expression [32, 46–48]. Given the fundamental role m^6^A plays in directing RNA fate and dynamics, targeting of the modification provides an opportunity to develop therapeutics which modulate the expression of genes essential for disease pathogenesis. During KSHV lytic reactivation, most cellular transcripts are degraded by a programme of host cell shutoff orchestrated by the viral SOX endonuclease [36, 37]. Therefore, genes that are resistant to the global pattern of downregulation may be more likely to be biologically relevant for KSHV lytic replication. As a result, we concentrated on m^6^A-modified transcripts with increased abundance in the lytic life cycle.

We identified four transcripts, *GPRC5A, STC1, FOSB* and *ZFP36L1* with increased mRNA levels during lytic replication; however, this effect could be modulated through the depletion of m^6^A machinery components, suggesting that m^6^A modification of these mRNAs contributes to their abundance. Accordingly, mutation of m^6^A sites within the *GPRC5A* transcript demonstrated that methylation increases its stability. Although m^6^A is generally thought to destabilise A/U base pairing, several studies have demonstrated that in certain sequence/structure-dependent contexts m^6^A improves mRNA stability, highlighting the importance of m^6^A examination on a site-specific basis [18, 22, 49]. Thus, we hypothesise that m^6^A allows increases in GPRC5A protein expression through tuneable improvements in mRNA stability, increasing the pool of mRNAs available for translation at any one time.

The importance of the m^6^A modified *GPRC5A* transcript in KSHV lytic replication is further reinforced via depletion studies, which show reduced lytic protein production and infectious virion production. Interestingly, GPRC5A was directly upregulated by ectopic expression of RTA, the master regulator of KSHV lytic replication. RTA binds the promoters of cellular and viral genes using its N-terminal DNA binding domain, while its C-terminal transactivation domain co-opts several host transcription factors to activate transcription [8, 10, 12]. Analysis of previous published RTA-ChIP-seq datasets revealed RTA-binding sites in the GPRC5A promoter, reinforcing our data that RTA upregulates these genes to play an important function within KSHV lytic replication [42]. Interestingly, GPRC5A was identified as one of many RTA-inducible cellular plasma membrane proteins which serve as key regulators of signalling and facilitate transformation of the host cell environment to support lytic replication [42].

As GPRC5A appears crucial to the process of lytic replication, we considered that identification of interaction partners might provide important insight into the role of GPRC5A within KSHV lytic replication. Quantitative mass spectrometry revealed that prominent interactors of GPRC5A were common constituents of plasma membranes, including the lipid raft structural proteins, namely members of the flotillin family, FLOT1 and FLOT2. We suggest this interaction signifies the organisation of GPRC5A, as with other GPCRs, into lipid rafts at the plasma membrane rather than providing any information as to the functional nature of GPRC5A within lytic reactivation. Interestingly however, the increased interaction of flotillins with GPRC5A during reactivation may suggest that GPRC5A undergoes greater packaging into lipid rafts in the lytic cycle to provide a microenvironment conducive to effective protein-protein interactions or signal transduction [50].

Previous research has identified that GPRC5A is a membrane protein, whose ligand is yet to be discovered, capable of activating cellular signalling pathways including STAT3 and EGFR [51–54] Although we tested whether GPRC5A was capable of modulating these pathways during KSHV reactivation by comparing scrambled control and GPRC5A-depleted TREx BCBL1-Rta cells, we were unable to detect EGFR activity in either latent or lytic replication states. Furthermore, STAT3 signalling, although rapidly inhibited during lytic replication, was unaffected by depletion of GPRC5A. Consequently, we focused on other key regulators of signalling, but no changes were also observed for pMEK1/2, pERK or pAKT. However, we identified an increase in levels of p-NFĸB S536 in cells depleted for GPRC5A, indicative of increased NFĸB signalling. Within cells latently infected with KSHV, NFκB is strongly activated by the viral latency factor vFLIP and drives the expression of latent transcripts while inhibiting lytic cycle-associated genes [45, 55]. Importantly, NFκB competes with RTA for binding of the Notch transcription factor RBP-JK and prevents transcription of RTA-responsive genes through this mechanism [45, 56]. Surprisingly however, our study and others have found that NFκB activity is consistently increased in the lytic cycle across a variety of cell lines [45]. This suggests that the inhibitory effect of NFκB on reactivation is at least partially negated or bypassed during lytic replication by additional cellular or viral factors, and results herein suggest that GPRC5A may modulate this complex relationship by restricting the activation of NFκB to levels which are insufficient to constrain KSHV lytic replication. However, the precise mechanism through which GPRC5A dampens NFκB activity during lytic replication remains unclear and further research is required to elucidate the intermediary factors which also play a role.

In summary, we show that m^6^A regulates the fate and function of cellular transcripts important for KSHV lytic replication. We identified the RTA-inducible gene GPRC5A, whose mRNA and protein abundance can be regulated by m^6^A-dependent alterations in RNA stability. Finally, we demonstrate that GPRC5A is important in regulating the inhibitory effect of NFκB signalling on KSHV lytic replication. Our work contributes further evidence to the fundamental nature of m^6^A in the posttranscriptional regulation of gene expression within viral infections. Given the ubiquitous distribution of RNA modifications across RNAs critical to viral infection, it seems likely that epitranscriptomics will emerge as a key determinant in the outcome of viral infections and a major contributor to the development of antiviral therapeutics.

## Materials and Methods

### Mammalian Cell culture

HEK 293T cells were obtained from the ATCC and were maintained in Dulbecco’s modified Eagle’s medium (DMEM) containing 10% foetal bovine serum (FBS) and 1% penicillin/streptomycin [57]. TREx BCBL1-Rta cells, a KSHV-infected, primary effusion lymphoma B lymphocyte cell line engineered to express C-myc-RTA under a doxycycline inducible promoter was kindly provided by Dr JU Jung, University of Southern California [58]. TREx BCBL1-Rta cells were cultured in RPMI 1640 medium (Sigma) with glutamine (Gibco) supplemented with 10% FBS, 1% P/S (Gibco) and 100 μg/ml hygromycin B (ThermoFisher). TREx BCBL1-Rta cells transduced with lentivirus were cultured in medium further supplemented with 3 μg/ml puromycin (ThermoFisher). KSHV lytic reactivation was induced in TREx BCBL1-Rta cells using 2 μg/ml doxycycline hyclate (Sigma). Cell lines were maintained at 37°C in a 5% CO_2_ atmosphere. Plasmids were transfected with Lipofectamine 2000 (ThermoFisher) in a 1:2 ratio in Opti-MEM media (Gibco) as previously described [59].

### Antibodies and plasmids

Antibodies used for immunoblotting: C-Myc (Sigma M4439 1:1000), FLOT1 (CST D2V7J 1:1000), FTO (Abcam ab126605 1:5000), GAPDH (Proteintech 60004-1-Ig 1:5000), GFP (Living Colours 632381 1:1000), GPRC5A (Atlas HPA007928 1:500), NFκB (CST D14E12), phospho-NFκB (CST 93H1), ORF57 (Santa Cruz sc-135747 1:1000), ORF65 (CRB crb2005224 1:100), STC1 (Proteintech 20621-1-AP 1:500), WTAP (Abcam ab195380 Rabbit 1:1000). pVSV.G and psPAX2 were gifted by Dr Edwin Chen (University of Westminster, London). GFP, GFP-ORF50, GFP-ORF57 plasmids are previously described [40, 41]. pRL-TK (Addgene #E2241) and pNFkB-ConA plasmids were gifted by the Macdonald group previously described in [60].The pLENTI-CMV-GFP-PURO construct was purchased from Addgene (#17448). GPRC5A, GPRC5A-FLAG and GPRC5A expressing plasmids were generated by PCR of the entire coding sequence of GPRC5A from TREx BCBL1-Rta cDNA and cloning using NEBuilder HIFI DNA assembly kit (NEB) into pLENTI-CMV-GFP-PURO. PLKO.1 TRC cloning vectors expressing shRNA sequences were purchased from Merck or Dharmacon: FTO (TRCN0000246247), GPRC5A KD1 (TRCN0000005628), GPRC5A KD2 (TRCN0000005632) WTAP (TRCN0000231423), YTHDF1 Sigma (TRCN0000286871). pLENTI-CMV-GPRC5A was mutated to remove m^6^A sites (A57G, G120T, C174T, A264G) using Q5 site directed mutagenesis kit (NEB) according to the manufacturer’s instructions. Primers are listed in **S3 Table**.

### RNA extraction, cDNA synthesis and qPCR

Total RNA was isolated from cells using the Monarch Total RNA Miniprep kit (NEB) according to the manufacturer’s protocol. 1 μg RNA was reverse transcribed to cDNA using LunaScript RT SuperMix Kit (NEB). qPCR was performed in 20 μl reactions containing diluted cDNA, GoTaq qPCR MasterMix (Promega) and the desired primer set. qPCR data was acquired using Rotorgene Q software and analysed using the ΔΔCT method relative to GAPDH as previously described [61].

### Immunoblotting

Protein samples were separated on 10–15% polyacrylamide gels and transferred to Amersham nitrocellulose membranes (GE healthcare) by Trans-blot Turbo Transfer system (Bio-Rad). Membranes were blocked in TBS + 0.1% Tween 20 with 5%(w/v) skimmed milk powder and probed with desired primary antibodies and secondary horseradish peroxidase conjugated IgG antibodies at 1:5000 (Dako Agilent). Protein bands were detected using a G:BOX (Syngene) after treating membranes with ECL Western Blotting Substrates (Promega). Densitometry was performed using ImageJ software.

### Immunofluorescence

TREx BCBL1-Rta cells were seeded onto coverslips treated with poly-L-lysine (Sigma), fixed for 15 mins in 4% paraformaldehyde and permeabilised with PBS + 1% Triton-X-100 as previously described [62]. Subsequent incubations were carried out at 37°C for 1h in a humidified chamber. Cells were blocked in PBS with 1% BSA prior to incubation with the desired primary and then Alexa-Fluor conjugated secondary antibodies. (Invitrogen 1:500). Coverslips were mounted onto slides using Vectashield Hardset Mounting Medium with DAPI (Vector laboratories). Images were acquired using a Zeiss LSM880 Inverted Confocal Microscope and analysed using ZEN 2009 imaging software (Carl Zeiss).

### Luciferase assays

HEK 293T cells were seeded for 24 hours before transfection with pNFkB-ConA (containing tandem kB-response element repeats)[63] and pRL-TK Renilla transfection control luciferase reporter plasmids. After 24 hours, samples were lysed with 1x Passive Lysis Buffer (Promega) and relative luciferase activity measured using Dual luciferase Stop and Glo reagents (Promega) as directed by the manufacturer.

### Viral reinfection assays

Virus-containing supernatant was harvested from TREx BCBL1-Rta cells reactivated for 72 h and used to infect naive HEK 293T cells in DMEM at a 1:1 ratio. 24 hours post-addition of viral supernatant, cells were collected, total RNA isolated and reverse transcribed into cDNA for qPCR analysis of ORF57 mRNA levels.

### Lentiviral transduction

Lentiviruses were produced by cotransfection of HEK 293T cells with 1.2 μg of lentiviral vector and 0.65 μg of packaging plasmids psPAX2 and VSV.G. After 72 hours, supernatant containing lentivirus was filter sterilised and combined with TREx BCBL1-Rta cells in 8 μg/ml polybrene (Merck Millipore). After 8 hours, the cells were placed in fresh medium. 48 hours post-transduction, cells were placed under selection with medium containing 3 μg/ml puromycin (Gibco) which was replaced every 2-3 days as previously described [64].

### m^6^A Immunoprecipitation

m^6^A IPs were carried out as described previously [32]. Briefly, TREx BCBL1-Rta cell total RNA was fragmented into 100-200 nucleotide segments using RNA fragmentation reagents (Ambion) and sodium acetate precipitated overnight at −80°C. A 5% input was collected prior to immunoprecipitation. The remaining RNA was combined with 25 μl of Magna ChIP Protein A+G magnetic beads (Merck Millipore) coated in 5μl of anti-m^6^A antibody (Merck Millipore) and incubated at 4°C overnight with rotation. RNA from inputs and m^6^A immunoprecipitations was eluted using proteinase K (ThermoFisher) followed by Trizol LS: chloroform extraction. RNA immunoprecipitations and inputs were converted to cDNA using LunaScript RT SuperMix Kit (NEB). m^6^A-immunoprecipitated samples were normalised to their respective input samples and m^6^A content at a particular region calculated relative to an unmodified control region within the same transcript.

### RNA stability assays

HEK 293T were transfected with appropriate plasmids for 48 hours then treated with 2.5 μg/ml of actinomycin D (Thermo Scientific) and samples collected at the desired time points. Total RNA was isolated prior to reverse transcription and qPCR analysis of specific transcripts as described previously. However, mRNA expression was normalised to 18s rRNA rather than GAPDH as the rRNA is more stable as previously described [32].

### Immunoprecipitation assays

GFP- or GPRC5A-GFP-transduced TREx BCBL1-Rta cells were lysed in GFP-TRAP buffer [10 mM Tris HCl pH7.5, 150 mM NaCl, 0.5 mM EDTA, 0.5% NP40 and incubated with 25 μl GFP-Trap Agarose beads (Chromotek) for 3 hours at 4°C. Beads were washed 3 times in GFP-TRAP buffer and immunoprecipitated proteins eluted by resuspension in 30 μl 2x Laemmli sample buffer and boiling at 90°C for 10 minutes. Finally, the proteins were separated by SDS-PAGE and detected by western blotting as described previously [65].

### Quantitative proteomics

Parental TREx BCBL1-Rta cells and GPRC5A-FLAG-expressing TREx BCBL1-Rta cells were lysed in immunoprecipitation buffer [150 mM NaCl, 2 mM EDTA, 0.5% Triton-X 100, 0.5 mM DTT, 10 mM Tris pH 7.4 and 1X protease and phosphatase inhibitors]. Lysates were centrifuged at 12,000 x g for 10 mins and the supernatant collected. A 5% input was taken at this stage and mixed with 15 μl 2x Laemmli sample buffer for downstream applications. 15 μl Protein A or G Dynabeads (ThermoFisher) were pre-prepared by washing 3 times in immunoprecipitation buffer then resuspended in 200μl immunoprecipitation buffer and coated with 2 μl FLAG M2 (Merck Millipore) or IgG (Merck Millipore, 12–370) control antibody. Beads were rotated for 1 hour at 4°C, washed 3 times with immunoprecipitation buffer and incubated with lysates overnight at 4°C. The following day, beads were washed 3 times in immunoprecipitation buffer and analysed at the University of Bristol Proteomics facility for tandem mass tagging (TMT) coupled with liquid chromatography and mass spectrometry analysis (LC-MS/MS) as previously described [66]. Data was outputted as a Microsoft Excel spreadsheet containing the results of a Sequest search against the Uniprot Human Database. Data from two independent replicates were grouped before filtering of best interactors. Contaminating proteins from a common contaminants database were removed and data filtered to satisfy an FDR of less than 5%. Protein abundance ratios were calculated between GPRC5A-FLAG cells and TREx BCBL1-Rta cells for both latently and lytically-infected cells. Proteins were filtered by a minimum abundance of 100 in GPRC5A-FLAG cells and a minimum net abundance of 50 after subtraction of background abundances in TREX cells. An abundance ratio of 2 was set for GPRC5A-FLAG cells relative to TREx BCBL1-Rta cells in latency. However, a 1.4-fold enrichment was used for GPRC5A-FLAG cells relative to TREx BCBL1-Rta cells in lytic replication as fewer proteins were detected due to the action of virus-mediated host-cell shutoff. Proteins with enriched GPRC5A interaction in the lytic phase were identified using a minimum 1.5-fold interaction in lytic cells compared to those undergoing latent replication after subtraction of TREx BCBL1-Rta cell background. The 51 proteins in latent, 51 proteins in lytic phase and 25 proteins enriched in the lytic phase which met these strict cut-offs were subjected to STRING protein-protein interaction networks functional enrichment analysis.

### Statistical analysis

Except otherwise stated, graphical data shown represent mean ± standard deviation of mean (SD) using three or more biologically independent experiments. Differences between means was analysed by unpaired Student’s t test calculated using Graphpad Prism 9 calculator. Statistics was considered significant at p < 0.05, with *P<0.05, **P<0.01, ***P<0.001.

### Publicly available deep-sequencing data

The following datasets were downloaded from the NCBI GEO database as raw FASTQ files: TREx BCBL1-RTA ChIP-seq, 4 input replicates (accessions: SRR8324529, SRR8324530, SRR8324531, SRR8324532) and 4 ChIP replicates (accessions: SRR8324533, SRR8324534, SRR8324535, SRR8324536). TREx BCBL1-RTA m^6^A IP, 2 input and IP replicates at KSHV infection timepoints of 0h (accessions: SRR7751622, SRR7751625, SRR7751628, SRR7751631); 8h (accessions: SRR7751623, SRR7751626, SRR7751629, SRR7751632); 24h (accessions: SRR7751624, SRR7751627, SRR7751630, SRR7751633). TREx BCBL1-RTA RNA-seq, 2 replicates at KSHV infection timepoints of 0h (accessions: SRR7751659, SRR7751663); 24h (accessions: SRR7751660, SRR7751664).

### ChIP-seq data processing

TREx BCBL1-RTA ChIP-seq datasets were subjected to quality control using FastQC (v0.11.9). The reads were mapped to the reference human genome GRCh38 using bwa (v0.7.17) with parameters mem –k 20 –T 15 –a –M –t 1. MACS2 (v2.2.7.1) was used to call peaks on corresponding input/CHIP bam files with parameters –g hs –B -- SPMR –q 0.01. The resulting output was used to generate fold change over input genome browser tracks for visualization using MACS2 bdgcmp –m FE.

### m^6^A-seq data processing

TREx BCBL1-RTA m^6^A-seq datasets were subjected to quality control using FastQC (v0.11.9) and mapped against a merger of the reference human GRCh38 and KSHV (NC_009333.1) genomes, as well as a junction database built from Ensembl 85, using STAR (v2.4.2a) with parameter --outFilterType BySJout. The resulting replicate bam files were collapsed into a single file using samtools (v1.9) merge and used to call peaks on corresponding input/IP bam files using MACS2 (v.2.2.7.1) with parameters –q 0.01 –nomodel–extsize 100 –B –SPMR –keep-dup all. The resulting output was used to generate fold change over input genome browser tracks for visualization using MACS2 bdgcmp –m FE. m^6^AViewer [67] called m^6^A sites used for calculating m^6^A changes were identified using the supplementary m^6^A_peak_calls table from GSE119026.

### RNA-seq data processing and analysis

TREx BCBL1-RTA RNA-seq datasets were adapter trimmed using Cutadapt (v1.9.1) with parameters –e 0.05 –minimum-length 15 and subjected to quality control and mapping as above. Read counts were extracted using featureCounts (v2.0.1) with parameters -M -O -C -B -p from a geneset of all merged gene trancripts from the Ensembl 85 annotation. The DESeq2 R package was then used to normalise the counts and calculate the relative changes in gene expression between latent and lytic conditions.

### Protein Pathway Analysis

Data were analyzed using Ingenuity Pathways Analysis (Ingenuity® Systems, www.ingenuity.com). Networks were generated using data sets containing gene identifiers and corresponding expression values that were uploaded into the application. Each gene identifier was mapped to its corresponding gene object in the Ingenuity Pathways Knowledge Base.

## Data availability

The mass spectrometry proteomics data have been deposited to the ProteomeXchange Consortium via the PRIDE partner repository. TREx BCBL1-RTA GPRC5A-FLAG Immunoprecipitation LC-MS. Project accession: PXD039669.

## Acknowledgments

We are grateful to members of the Whitehouse laboratory for helpful discussions. We thank Professor Jae Jung (University of Southern California School of Medicine, Los Angeles) for the TREx BCBL1-RTA cells, Dr Edwin Chen (University of Westminster) for the lentivirus vectors and Dr. Kate Heesom (Proteomics Facility, University of Bristol, UK) for proteomic technical assistance and initial sorting of proteomic data. The work was supported in parts by the Biotechnology and Biological Sciences Research Council (BBSRC) (BB/M006557/1) and Medical Research Council (MR/R010145/1, MR/X000060/1) project grants and a MRC DiMEN DTP grant (95505168). IDY and SAW acknowledge support from BBSRC (BB/V00722X/1).

## Supporting Information Legends

**S1 Fig. Differential m^6^A modification of cellular transcripts during KSHV lytic replication.** Genome sequencing tracks for (A) *GPRC5A*, (B) *FOSB*, (C) *STC1* and (D) *ZFP36L1* from latent and lytic TREx BCBL1‐Rta cells depicting the m^6^A IP fold enrichment over input on cellular mRNAs. Differentially modified peaks showing an increase in m^6^A modification during lytic replication are indicated by black boxes.

**S2 Fig. Canonical pathways, molecular and cellular functions and diseases associated with differentially methulated cellular m^6^A mRNAs.** Ingenuity pathway analysis (IPA) was performed to predict associated canonical pathways, molecular and cellular functions, and diseases associated with the differentially methylated m^6^A mRNAs during KSHV lytic replication.

**S3 Fig. Mutation of DRACH sequences within GPRC5A to abolish the differentially modified m^6^A peak during KSHV lytic replication.** Site directed mutagenesis was carried out to abrogate 4 DRACH sequences of GPRC5A in close proximity to the m^6^A peak identified as differentially modified during KSHV lytic replication. Consensus DRACH sequences are highlighted with red box.

**S4 Fig. RTA binds the promoters and internally within GPRC5A and ZFP36L1.** Genome sequencing tracks from lytic TREx BCBL1‐Rta cells depicting RTA‐CHIP peaks within (A) GPRC5A and (B) ZFP36L1 genes. Experiments were performed in Papp et al 2019 [42].

**S1 Table. List of transcripts with differentially modified m^6^A peaks during KSHV lytic replication.** Differentially modified m^6^A peaks were determined relative to stable m^6^A peaks within the same transcript during KSHV lytic replication. Raw and relativized to 0h peak intensity values are listed for the differentially modified m^6^A peak within each cellular transcript. Sites where m^6^A peaks were called significant by m^6^A viewer software are highlighted in blue. The overall change in methylation is also included. The DESeq2 calculated changes in gene expression of the selected transcripts are indicated as well as the overall change in methylation and gene expression.

**S2 Table. List of host cell proteins with an enhanced interaction with GPRC5A during KSHV lytic replication.** Global quantitative proteomic analysis was performed on FLAG immunoprecipitates from latent and reactivated TREx BCBL1‐Rta cell lysates. Proteins with an enhanced interaction with GPRC5A during lytic replication are listed.

**S3 Table. List of primer sequences.**

