## Supplementary figs and tables for "m^6^A regulates the stability of cellular transcripts required for efficient KSHV lytic replication"

**Supporting Information**

**Supplementary Figures 1-4**
**Supplementary Tables 1-3**

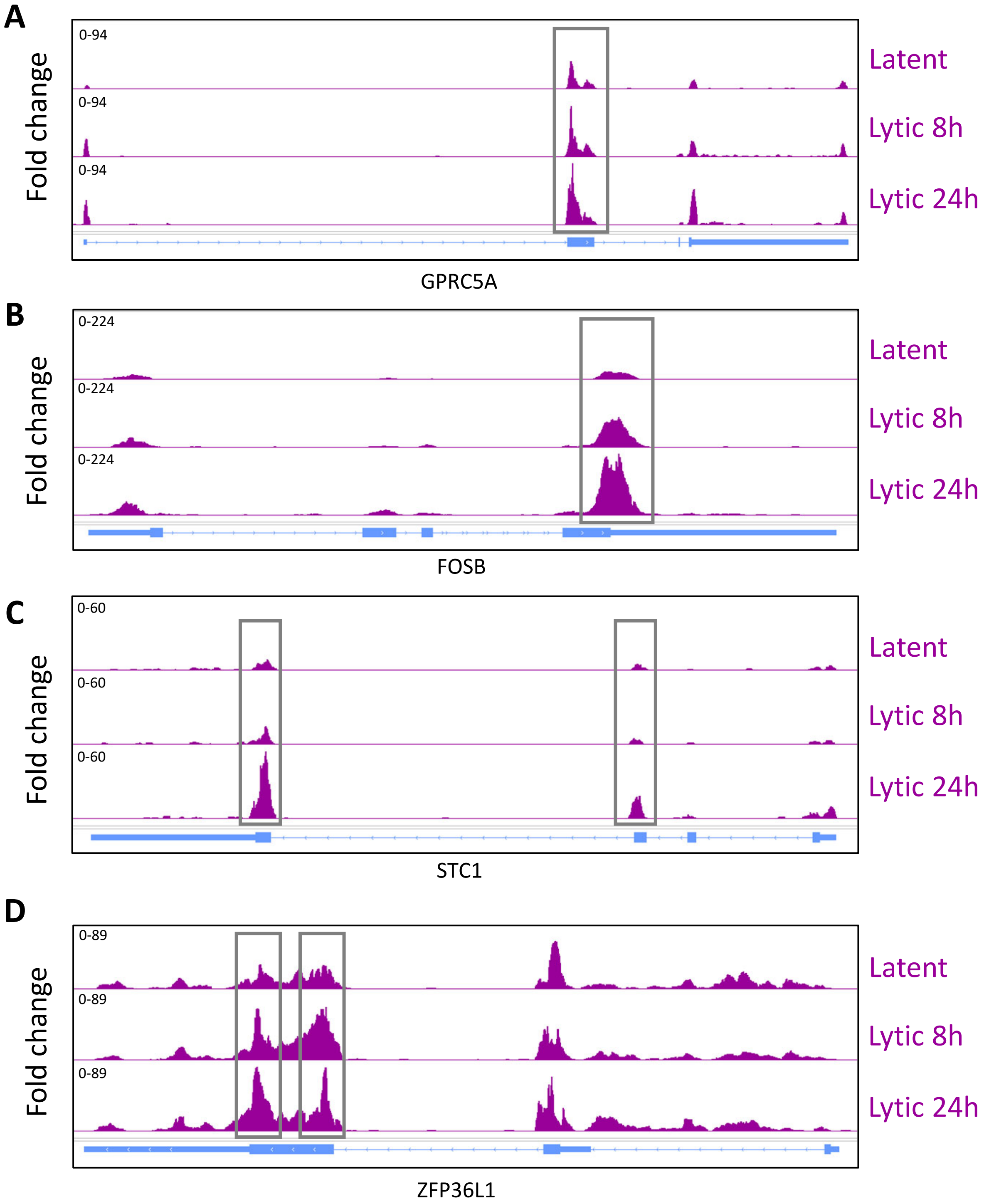
A
Fold change
0-94
0-94
0-94
Latent
Lytic 8h
Lytic 24h
GPRC5A
B
Fold change
0-224
0-224
0-224
Latent
Lytic 8h
Lytic 24h
FOSB
C
Fold change
0-60
0-60
0-60
Latent
Lytic 8h
Lytic 24h
STC1
D
Fold change
0-89
0-89
0-89
Latent
Lytic 8h
Lytic 24h
ZFP36L1

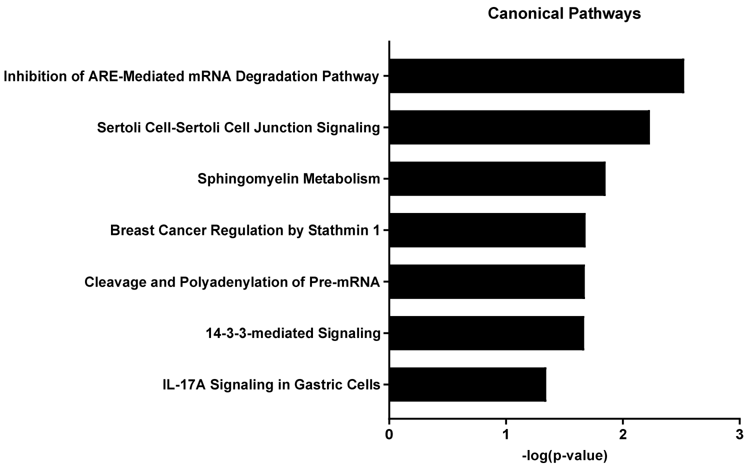
Canonical Pathways
Inhibition of ARE-Mediated mRNA Degradation Pathway
Sertoli Cell-Sertoli Cell Junction Signaling
Sphingomyelin Metabolism
Breast Cancer Regulation by Stathmin 1
Cleavage and Polyadenylation of Pre-mRNA
14-3-3-mediated Signaling
IL-17A Signaling in Gastric Cells
0
1
2
3
-log(p-value)

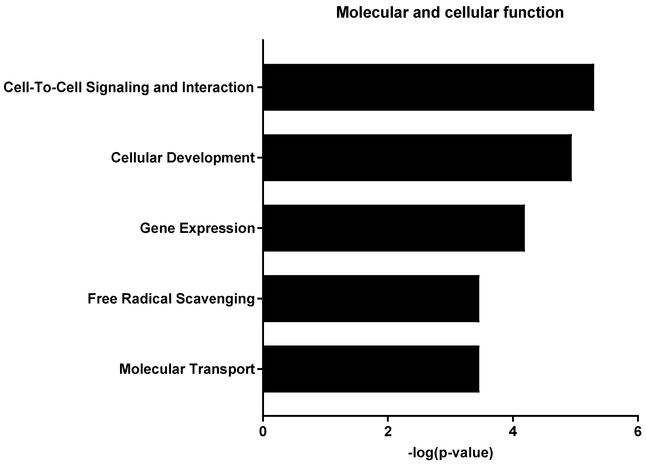
Molecular and cellular function
Cell-To-Cell Signaling and Interaction
Cellular Development
Gene Expression
Free Radical Scavenging
Molecular Transport
0
2
4
6
-log(p-value)

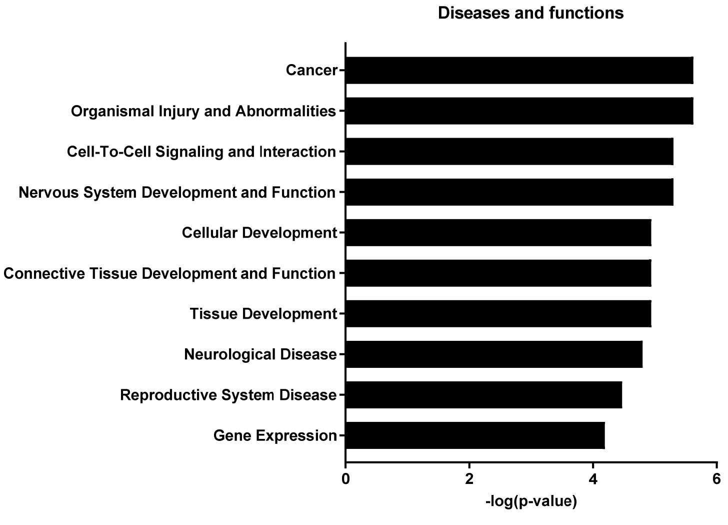
Diseases and functions
Cancer
Organismal Injury and Abnormalities
Cell-To-Cell Signaling and Interaction
Nervous System Development and Function
Cellular Development
Connective Tissue Development and Function
Tissue Development
Neurological Disease
Reproductive System Disease
Gene Expression
0
2
4
6
-log(p-value)

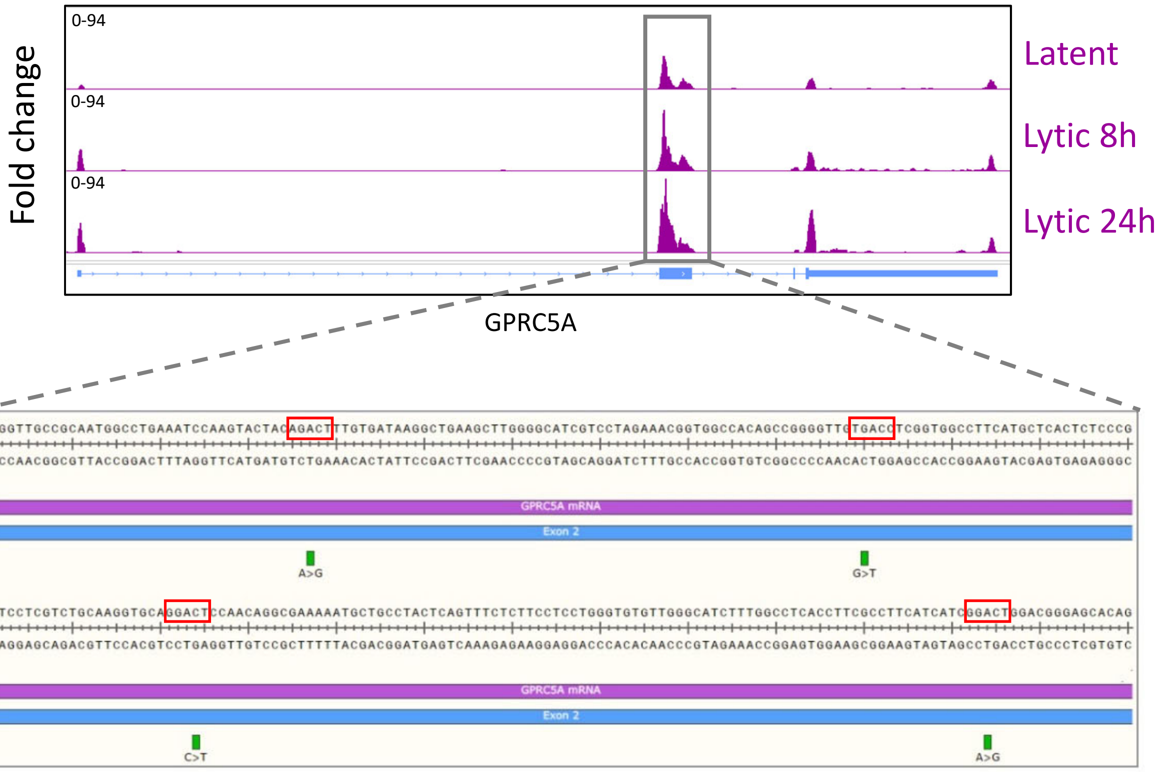

Fold change
0-94
0-94
0-94
Latent
Lytic 8h
Lytic 24h
GPRC5A
GPRC5A mRNA
Exon 2
A>G
G>T
GPRC5A mRNA
Exon 2
C>T
A>G

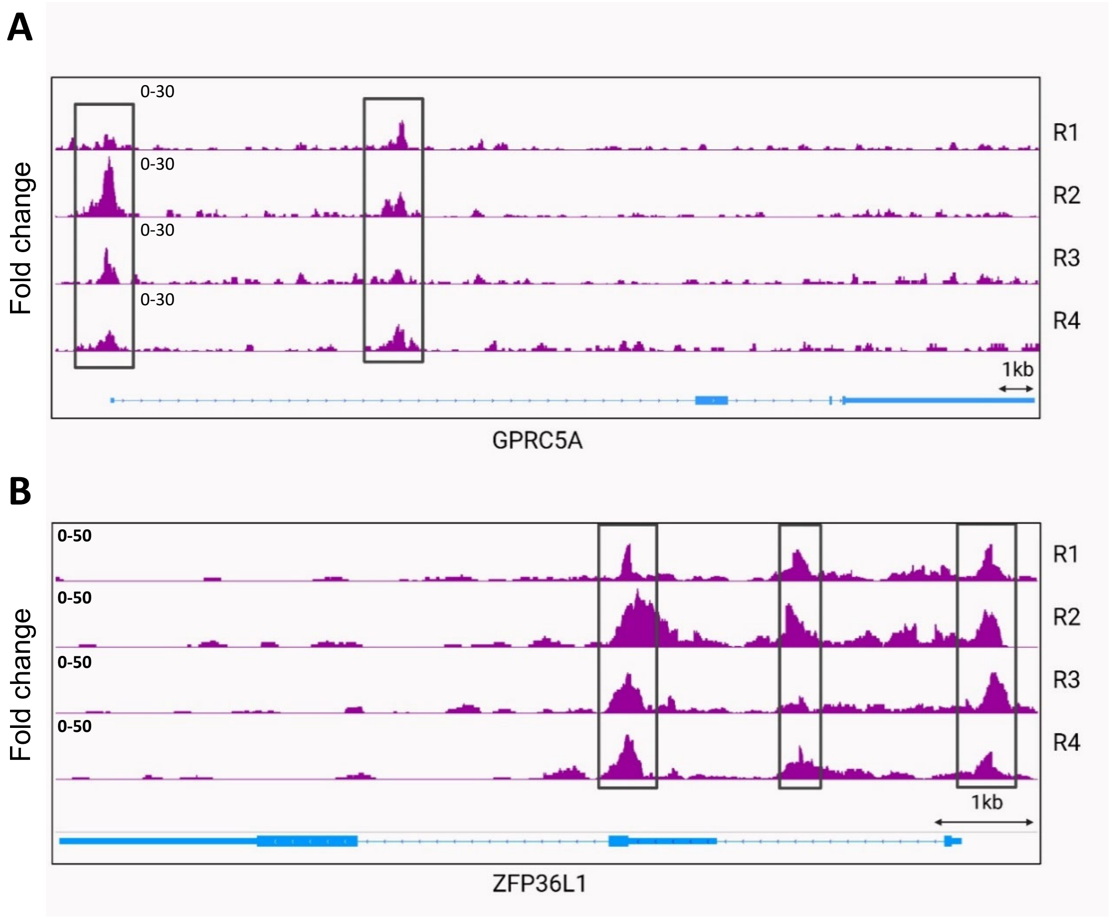
A
0-30
0-30
0-30
0-30
R1
R2
R3
R4
Fold change
1kb
GPRC5A
B
0-50
0-50
0-50
0-50
R1
R2
R3
R4
Fold change
1kb
ZFP36L1

Dark grey = low expression

| Gene | Raw values (IGV) R1 0h | Raw values (IGV) R1 8h | Raw values (IGV) R1 20h | Raw values (IGV) R2 0h | Raw values (IGV) R2 8h | Raw values (IGV) R2 20h | Relative values (IGV) R1 0h | Relative values (IGV) R1 8h | Relative values (IGV) R1 20h | Relative values (IGV) R2 0h | Relative values (IGV) R2 8h | Relative values (IGV) R2 20h | Relative expression change (DESeq2) R1+R2 0h | Relative expression change (DESeq2) R1+R2 20h | Methlyation change | Expression change |
|---|---|---|---|---|---|---|---|---|---|---|---|---|---|---|---|---|
| ADIRF | 0.16 | 0.08 | 0.05 | 1.67 | 0.67 | 0.09 | 1.00 | 0.50 | 0.31 | 1.00 | 0.40 | 0.05 | | | Decrease | |
| ADGRA1 | 0.04 | 0.08 | 0.06 | 0.05 | 0.16 | 0.16 | 1.00 | 2.00 | 1.50 | 1.00 | 3.20 | 3.20 | | | Increase | |
| ATF3 | 0.04 | 0.04 | 0.32 | 0.05 | 0.10 | 0.19 | 1.00 | 1.00 | 8.00 | 1.00 | 2.00 | 3.80 | 1.00 | 25.67 | Increase | Increase |
| BAIAP2 | 0.02 | 0.03 | 0.17 | 0.12 | 0.19 | 0.21 | 1.00 | 1.50 | 8.50 | 1.00 | 1.58 | 1.75 | 1.00 | 51.12 | Increase | Increase |
| BAMBI | 1.39 | 1.04 | 1.58 | 5.76 | 7.04 | 1.49 | 1.00 | 0.75 | 1.14 | 1.00 | 1.22 | 0.26 | 1.00 | 3.53 | Decrease | Increase |
| BRD9 | 1.02 | 0.82 | 0.89 | 16.00 | 10.00 | 2.14 | 1.00 | 0.80 | 0.87 | 1.00 | 0.63 | 0.13 | 1.00 | 0.94 | Decrease | Decrease |
| CHRNA3 | 0.02 | 0.04 | 0.38 | 0.04 | 0.25 | 0.29 | 1.00 | 2.00 | 19.00 | 1.00 | 6.25 | 7.25 | 1.00 | 4.23 | Increase | Increase |
| CNOT2 | 0.24 | 0.21 | 0.21 | 2.76 | 2.30 | 0.24 | 1.00 | 0.88 | 0.88 | 1.00 | 0.83 | 0.09 | 1.00 | 0.68 | Decrease | Decrease |
| COL23A1 | 0.08 | 0.12 | 0.05 | 0.30 | 0.24 | 0.09 | 1.00 | 1.50 | 0.63 | 1.00 | 0.80 | 0.30 | 1.00 | 9.64 | Decrease | Increase |
| DLG2 | 0.01 | 0.01 | 0.05 | 0.05 | 0.04 | 0.02 | 1.00 | 1.00 | 5.00 | 1.00 | 0.87 | 0.43 | 1.00 | 3.07 | Decrease | Increase |
| DPYSL2 | 0.05 | 0.02 | 0.34 | 0.03 | 0.31 | 0.28 | 1.00 | 0.40 | 6.80 | 1.00 | 10.33 | 9.33 | 1.00 | 2.23 | Increase | Increase |
| EPPK1 | 0.33 | 0.23 | 0.23 | 1.93 | 1.87 | 0.72 | 1.00 | 0.70 | 0.70 | 1.00 | 0.97 | 0.37 | 1.00 | 9.18 | Decrease | Increase |
| EPS8 | 0.05 | 0.04 | 0.11 | 0.07 | 0.20 | 0.20 | 1.00 | 0.80 | 2.20 | 1.00 | 2.86 | 2.86 | 1.00 | 3.18 | Increase | Increase |
| FOSB | 0.06 | 0.12 | 1.74 | 0.63 | 3.02 | 1.94 | 1.00 | 2.00 | 29.00 | 1.00 | 4.79 | 3.08 | 1.00 | 16.15 | Increase | Increase |
| GALNT9 | 0.01 | 0.04 | 0.04 | 0.05 | 0.06 | 0.06 | 1.00 | 4.00 | 4.00 | 1.00 | 1.20 | 1.20 | 1.00 | 7.83 | Increase | Increase |
| GOLM1 | 0.22 | 0.20 | 0.09 | 1.06 | 1.91 | 0.09 | 1.00 | 0.91 | 0.41 | 1.00 | 1.80 | 0.08 | 1.00 | 1.47 | Decrease | Increase |
| GPRC5A | 0.07 | 0.07 | 0.70 | 0.88 | 2.79 | 1.18 | 1.00 | 1.00 | 10.00 | 1.00 | 3.17 | 1.34 | 1.00 | 18.49 | Increase | Increase |
| HEY1 | 0.01 | 0.03 | 0.22 | 0.03 | 0.80 | 0.22 | 1.00 | 3.00 | 22.00 | 1.00 | 26.67 | 7.33 | 1.00 | 109.04 | Increase | Increase |
| hnRNPA3 | 0.31 | 0.21 | 0.04 | 3.07 | 3.31 | 0.15 | 1.00 | 0.68 | 0.13 | 1.00 | 1.08 | 0.05 | 1.00 | 0.50 | Decrease | Decrease |
| IL12RB2 | 0.01 | 0.01 | 0.36 | 0.04 | 0.08 | 0.06 | 1.00 | 1.00 | 36.00 | 1.00 | 2.00 | 1.50 | 1.00 | 5.43 | Increase | Increase |
| ILF3 | 0.04 | 0.07 | 0.47 | 0.10 | 2.08 | 0.73 | 1.00 | 1.75 | 11.75 | 1.00 | 20.80 | 7.30 | 1.00 | 0.93 | Decrease | Decrease |
| JTB | 0.09 | 0.05 | 0.02 | 1.07 | 0.16 | 0.04 | 1.00 | 0.56 | 0.22 | 1.00 | 0.15 | 0.04 | 1.00 | 0.96 | Decrease | Decrease |
| JUN | 0.45 | 0.39 | 6.46 | 5.41 | 19.00 | 3.64 | 1.00 | 0.87 | 14.36 | 1.00 | 3.51 | 0.67 | 1.00 | 4.74 | Increase | Increase |
| MAGEC1 | 0.10 | 0.10 | 0.05 | 1.14 | 0.65 | 0.07 | 1.00 | 1.00 | 0.50 | 1.00 | 0.57 | 0.06 | 1.00 | 0.45 | Decrease | Decrease |
| MCF2L | 0.01 | 0.07 | 0.12 | 0.37 | 0.44 | 0.35 | 1.00 | 7.00 | 12.00 | 1.00 | 1.19 | 0.95 | 1.00 | 5.79 | Increase | Increase |
| PABPN1 | 0.47 | 0.33 | 0.44 | 1.85 | 1.52 | 0.17 | 1.00 | 0.70 | 0.94 | 1.00 | 0.82 | 0.09 | 1.00 | 0.68 | Decrease | Decrease |
| PCF11 | 0.44 | 0.28 | 0.37 | 2.74 | 2.77 | 0.38 | 1.00 | 0.64 | 0.84 | 1.00 | 1.01 | 0.14 | 1.00 | 1.49 | Decrease | Increase |
| PCNX1 | 0.04 | 0.04 | 0.60 | 0.62 | 1.16 | 1.16 | 1.00 | 1.00 | 15.00 | 1.00 | 1.87 | 1.87 | 1.00 | 0.82 | Increase | Decrease |
| PEAR1 | 0.01 | 0.03 | 0.06 | 0.08 | 0.32 | 0.17 | 1.00 | 3.00 | 6.00 | 1.00 | 4.00 | 2.13 | 1.00 | 3.14 | Increase | Increase |
| PILRB | 0.06 | 0.04 | 0.48 | 0.16 | 1.03 | 0.63 | 1.00 | 0.67 | 8.00 | 1.00 | 6.44 | 3.94 | 1.00 | 1.73 | Increase | Increase |
| PLEKHA6 | 0.02 | 0.05 | 0.30 | 0.05 | 0.38 | 0.47 | 1.00 | 2.50 | 15.00 | 1.00 | 7.60 | 9.40 | 1.00 | 2.70 | Increase | Increase |
| PPP2R2D | 1.01 | 0.58 | 0.77 | 5.18 | 7.96 | 1.18 | 1.00 | 0.57 | 0.76 | 1.00 | 1.54 | 0.23 | 1.00 | 0.90 | Decrease | Decrease |
| PRDM16 | 0.03 | 0.05 | 0.11 | 0.01 | 0.17 | 0.22 | 1.00 | 1.67 | 3.67 | 1.00 | 17.00 | 22.00 | 1.00 | 9.91 | Increase | Increase |
| PTPRN2 | 0.01 | 0.04 | 0.10 | 0.03 | 0.48 | 0.59 | 1.00 | 4.00 | 10.00 | 1.00 | 16.00 | 19.67 | 1.00 | 4.33 | Increase | Increase |
| RBM12B | 4.13 | 3.07 | 0.90 | 58.00 | 32.00 | 4.64 | 1.00 | 0.74 | 0.22 | 1.00 | 0.55 | 0.08 | 1.00 | 1.10 | Decrease | Increase |
| RFTN1 | 0.30 | 0.27 | 0.02 | 1.68 | 0.99 | 0.12 | 1.00 | 0.90 | 0.07 | 1.00 | 0.59 | 0.07 | 1.00 | 0.63 | Decrease | Decrease |
| RhoU | 0.24 | 0.19 | 0.92 | 2.08 | 4.37 | 3.81 | 1.00 | 0.79 | 3.83 | 1.00 | 2.10 | 1.83 | 1.00 | 0.25 | Increase | Decrease |
| SDF4 | 0.02 | 0.06 | 0.33 | 0.11 | 0.50 | 0.42 | 1.00 | 3.00 | 16.50 | 1.00 | 4.55 | 3.82 | 1.00 | 1.55 | Increase | Increase |
| SGMS2 | 0.06 | 0.11 | 0.89 | 0.37 | 5.77 | 1.13 | 1.00 | 1.83 | 14.83 | 1.00 | 15.59 | 3.05 | 1.00 | 5.67 | Increase | Increase |
| SLC25A25 | 0.06 | 0.05 | 0.26 | 0.09 | 1.21 | 0.12 | 1.00 | 0.83 | 4.33 | 1.00 | 13.44 | 1.33 | 1.00 | 3.25 | Increase | Increase |
| SLC6A9 | 0.03 | 0.05 | 0.32 | 0.08 | 1.45 | 0.27 | 1.00 | 1.67 | 10.67 | 1.00 | 18.13 | 3.38 | 1.00 | 2.97 | Increase | Increase |
| STC1 | 0.03 | 0.09 | 0.52 | 0.21 | 0.40 | 0.45 | 1.00 | 3.00 | 17.33 | 1.00 | 1.90 | 2.14 | 1.00 | 4.54 | Increase | Increase |
| TEX14 | 0.01 | 0.03 | 0.09 | 0.06 | 0.03 | 0.13 | 1.00 | 3.00 | 9.00 | 1.00 | 0.50 | 2.17 | 1.00 | 5.88 | Increase | Increase |
| TIFA | 0.76 | 0.38 | 0.09 | 5.91 | 3.84 | 0.40 | 1.00 | 0.50 | 0.12 | 1.00 | 0.65 | 0.07 | 1.00 | 0.97 | Decrease | Decrease |
| TNN | 0.32 | 0.38 | 1.34 | 5.63 | 8.15 | 3.74 | 1.00 | 1.19 | 4.19 | 1.00 | 1.45 | 0.66 | 1.00 | 1.56 | Decrease | Increase |
| TNRC6A | 1.33 | 1.04 | 2.34 | 9.47 | 18.00 | 2.35 | 1.00 | 0.78 | 1.76 | 1.00 | 1.90 | 0.25 | 1.00 | 0.72 | Decrease | Decrease |
| TUBB3 | 0.04 | 0.04 | 0.04 | 0.18 | 0.20 | 0.08 | 1.00 | 1.00 | 1.00 | 1.00 | 1.11 | 0.44 | 1.00 | 1.55 | Decrease | Increase |
| UAP56 | 0.01 | 0.01 | 0.62 | 0.02 | 0.33 | 0.78 | 1.00 | 1.00 | 62.00 | 1.00 | 16.50 | 39.00 | 1.00 | 2.01 | Increase | Increase |
| USP48 | 0.07 | 0.08 | 0.21 | 0.68 | 1.12 | 0.18 | 1.00 | 1.14 | 3.00 | 1.00 | 1.65 | 0.26 | 1.00 | 0.79 | Decrease | Decrease |
| YBEY | 0.21 | 0.18 | 0.09 | 2.53 | 1.32 | 0.15 | 1.00 | 0.86 | 0.43 | 1.00 | 0.52 | 0.06 | 1.00 | 0.93 | Decrease | Decrease |
| YTHDF3 | 1.72 | 1.57 | 2.07 | 12.00 | 23.00 | 3.64 | 1.00 | 0.91 | 1.20 | 1.00 | 1.92 | 0.30 | 1.00 | 0.76 | Decrease | Decrease |
| ZBTB21 | 0.03 | 0.14 | 0.31 | 0.13 | 1.17 | 0.59 | 1.00 | 4.67 | 10.33 | 1.00 | 9.00 | 4.54 | 1.00 | 1.19 | Increase | Increase |
| ZC3H12A | 0.24 | 0.08 | 0.13 | 1.06 | 1.19 | 0.25 | 1.00 | 0.33 | 0.54 | 1.00 | 1.12 | 0.24 | 1.00 | 2.75 | Decrease | Increase |
| ZFP36L1 | 0.17 | 0.19 | 2.55 | 1.66 | 3.55 | 2.46 | 1.00 | 1.12 | 15.00 | 1.00 | 2.14 | 1.48 | 1.00 | 4.54 | Increase | Increase |
| ZNF12 | 0.20 | 0.18 | 0.35 | 13.00 | 3.16 | 0.35 | 1.00 | 0.90 | 1.75 | 1.00 | 0.24 | 0.03 | 1.00 | 0.99 | Decrease | Decrease |

| Gene names | Lytic/Latent ratio |
|---|---|
| FLOT2 | 26.72925764 |
| SRRM1 | 6.875840979 |
| POLR1G | 4.852941176 |
| ACTL8 | 4.740740741 |
| FLOT1 | 4.205438066 |
| SCAF4 | 3.354297694 |
| LUC7L3 | 3.269565217 |
| PINX1 | 2.61774744 |
| WDR6 | 2.212 |
| EWSR1 | 1.787741203 |
| ARC | 1.782608696 |
| CD44 | Interaction exclusive to lytic replication |
| FUBP1 | Interaction exclusive to lytic replication |
| PPIF | Interaction exclusive to lytic replication |
| VDAC1 | Interaction exclusive to lytic replication |
| VDAC2 | Interaction exclusive to lytic replication |
| VDAC3 | Interaction exclusive to lytic replication |

| Gene name | Forward | Reverse |
|---|---|---|
| FOSB | 5'-GAAATGCCCGGTTCCTTC-3' | 5'-GAGGGTGGGTTGCACAAG-3' |
| FTO | 5'-TCTGACCCCCAAAGATGATG-3' | 5'-CTCGGAGAATTAGTTTAGGATATTTCA-3' |
| GAPDH | 5'-TGTCAGTGGTGGACCTGAC-3' | 5'-GTGGTCGTTGAGGGCAATG-3' |
| GPRC5A | 5'-CCTTTCCCTGTTGGTGATTCT -3' | 5'-AGACATTGACGTTGGTCCTATTC-3' |
| JUN | 5'-CGCCTGATAATCCAGTCCA-3' | 5'-TTCTTGGGGCACAGGAACT-3' |
| ORF47 | 5'-CGCGGTCGTTCGAAGATTG-3' | 5'-CGAGTCTGACTTCCGCTAA-3' |
| ORF57 | 5'-GCCATAATCAAGCGTACTGG-3' | 5'-GCAGACAAATATTGCGGTGT-3' |
| PLEKHA6 | 5'-CCACCAGGCAAGAGGTAGAG-3' | 5'-GCACAACGCCAACTTTGTT-3' |
| STC1 | 5'-AGGCGGAGCAGAATGACTC-3' | 5'-GTTGAGGCAACGAACCACTT-3' |
| WTAP | 5'-TTCCCAAGAAGGTTCGATTG-3' | 5'-TGCAGACTCCTGCTGTTGTT-3' |
| YTHDF1 | 5'-ATAACCAGCTCCGGCACAT-3' | 5'-GGGAGTTTGTGACCGGTTT-3' |
| ZC3H12A | 5'-TCTGTGGGAATTTGAGGACAG-3' | 5'-GTGGATCTCCGTGGATGAATAG-3' |
| ZFP36L1 | 5'-GCGAAGTTTTATGCAAGGGTAA-3' | 5'-GTGCCCACTGCCTTTCTG-3' |
| FOSB Control | 5'-GGAGAGCTGGTGACTTTGGG-3' | 5'-AGAGCCAACAGTCAGCTGGG-3' |
| FOSB m$^6$A | 5'-CTCCCTCCTCGCTCTGTGAA-3' | 5'-CAAGTCTCTCTCCCCCATGT-3' |
| GAPDH 1 | 5'-GCATCTTCTTTTGCGTCGCC-3' | 5'-TTGACTCCGACCTTCACCTTCC-3' |
| GAPDH 2 | 5'-TGCACCACCAACTGCTTA-3' | 5'-ATGAGTCCTTCCACGATACC-3' |
| GPRC5A control | 5'-CCGAGATCTAATCTCCCCCTA-3' | 5'-GGGCTTGTGCTAGTGAGGTC-3' |
| GPRC5A m$^6$A | 5'-CTCACTCTCCCGATCCTCGT-3' | 5'-GAAACTGAGTAGGCAGCATT-3' |
| JUN Ctrl | 5'-AGCGCCTGATAATCCAGTCC-3' | 5'-ATCTGTCACGTTCTTGGGGC-3' |
| JUN m$^6$A | 5'-ACCTTGAAAGCTCAGAACTCGG-3' | 5'-TAAGCTGTGCCACCTGTTCC-3' |
| SLC39A14 control | 5'-GCAGGATCTAATACATCGGTATGG-3' | 5'-TGGTTGAGTAGGGCCTTCAG-3' |
| SLC39A14 m$^6$A | 5'-GGACAGATCCAGATTGGGTAG-3' | 5'-AGGGCCCGACTTCCAGT-3' |
| ZFP36L1 control | 5'-GGCACACACACATTAAGATGAA-3' | 5'-AGAGAAATAGAAAGCGACGGT-3' |
| ZFP36L1 m$^6$A site 1 | 5'-GGTTGCCTGCTGGACAGAAA -3' | 5'-TTCTGGTGGAACTTGGAGCTG-3' |
| ZFP36L1 m$^6$A site 2 | 5'-CAGGATTCTCTCTCGGACCA -3' | 5'-TCCAAGGTCGGGGAGTCT-3' |
| WTAP shRNA KD | TRCN0000231423 | |
| FTO shRNA KD | TRCN0000246247 | |
| YTHDF1 shRNA KD | TRCN0000286871 | |
| GPRC5A shRNA KD1 | TRCN0000005628 | |
| GPRC5A shRNA KD2 | TRCN0000005632 | |
